# Vaccination with mRNA-encoded membrane-bound HIV Envelope trimer induces neutralizing antibodies in animal models

**DOI:** 10.1101/2025.01.24.634423

**Authors:** Parham Ramezani-Rad, Christopher A. Cottrell, Ester Marina-Zárate, Alessia Liguori, Elise Landais, Jonathan L. Torres, Amber Myers, Jeong Hyun Lee, Sabyasachi Baboo, Claudia Flynn, Katherine McKenney, Eugenia Salcedo, Xiaoya Zhou, Oleksandr Kalyuzhniy, Erik Georgeson, Nicole Phelps, Danny Lu, Saman Eskandarzadeh, Sergey Menis, Michael Kubitz, Bettina Groschel, Nushin Alavi, Abigail M. Jackson, Wen-Hsin Lee, Andy S. Tran, Elana Ben-Akiva, Katarzyna Kaczmarek Michaels, Jolene K. Diedrich, Chiamaka A. Enemuo, Vanessa Lewis, Arpan Pradhan, Sudhir Pai Kasturi, Torben Schiffner, Jon M. Steichen, Diane G. Carnathan, Sunny Himansu, John R. Yates, James C. Paulson, Gabriel Ozorowski, Darrell J. Irvine, Guido Silvestri, Devin Sok, Andrew B. Ward, Shane Crotty, William R. Schief

**Affiliations:** Center for Vaccine Innovation, La Jolla Institute for Immunology, La Jolla, CA 92037, USA; Consortium for HIV/AIDS Vaccine Development (CHAVD), The Scripps Research Institute, La Jolla, CA 92037, USA; Department of Immunology and Microbiology, The Scripps Research Institute, La Jolla, CA 92037, USA; IAVI Neutralizing Antibody Center, San Diego, CA 92121, USA; Department of Integrative Structural and Computational Biology, The Scripps Research Institute, La Jolla, CA 92037, USA; Department of Molecular Medicine, The Scripps Research Institute, La Jolla, CA 92037, USA; Department of Biological Engineering, Massachusetts Institute of Technology, Cambridge, MA 02139, USA; Emory National Primate Research Center and Emory Vaccine Center, Emory University School of Medicine, Atlanta, GA 30329, USA; Moderna, Inc. Cambridge, MA 02139, USA; Howard Hughes Medical Institute, Chevy Chase, MD 20815, USA; Department of Medicine, Division of Infectious Diseases and Global Public Health, University of California, San Diego (UCSD), La Jolla, CA 92037, USA

## Abstract

A protective vaccine against HIV will likely need to induce broadly neutralizing antibodies (bnAbs) that engage relatively conserved epitopes on the HIV envelope glycoprotein (Env) trimer. Nearly all vaccine strategies to induce bnAbs require the use of relatively complex immunization regimens involving a series of different immunogens, most of which are Env trimers. Producing protein-based clinical material to evaluate such relatively complex regimens in humans presents major challenges in cost and time. Furthermore, immunization with HIV trimers as soluble proteins induces strong non-neutralizing responses to the trimer base, which is normally occluded on the virion. These base responses could potentially detract from the induction of nAbs and the eventual induction of bnAbs. mRNA vaccine platforms offer potential advantages over protein delivery for HIV vaccine development, including increased production speed, reduced cost, and the ability to deliver membrane-bound trimers that might facilitate improved immuno-focusing to non-base epitopes. We report the design of mRNA-delivered soluble and membrane-bound forms of a stabilized native-like Env trimer (BG505 MD39.3), initial immunogenicity evaluation in rabbits that triggered clinical evaluation, and more comprehensive evaluation of B cell, T cell, and antibody responses in non-human primates. mRNA-encoded membrane-bound Env immunization elicited reduced off-target base-directed Env responses and stronger neutralizing antibody responses, compared with mRNA-encoded soluble Env. Overall, mRNA delivery of membrane-bound Env appears promising for enhancing B cell responses to subdominant epitopes and facilitating rapid translation to clinical testing, which should assist HIV vaccine development.

**One Sentence Summary:** HIV envelope trimer mRNA enables membrane-bound expression and represents a functional immunogen in pre-clinical mammalian models.

## Introduction

Effective vaccine strategies are required to build immunity to pathogens with significant health threats. The HIV pandemic has been recognized since 1981, but HIV continues to infect and kill millions of people annually (*1*). HIV represents a remarkable vaccine challenge due to its genetic diversity, immune escape and integration into the host genome (*2*). The Env trimer on the surface of HIV is the sole target of neutralizing antibodies (nAbs); and nAbs to different Env epitopes can confer protective responses to varying amounts of HIV strains (*3*). However, nAbs are difficult to elicit due to the immunodominance of non-neutralizing epitopes on Env immunogens and the abundant glycosylation of the Env trimer that limits accessibility to neutralizing epitopes. Owing to the antigenic diversity of HIV Env across different isolates, most vaccine design strategies aim to induce broadly neutralizing antibodies (bnAbs), defined as nAbs that neutralize diverse HIV isolates (*2*). Inducing bnAbs faces additional challenges of the low frequency of naive bnAb-precursor B cells having genetic and structural potential to develop into bnAbs and the need to guide somatic hypermutation (SHM) to produce bnAbs. Nearly all bnAb vaccine strategies currently being pursued involve sequential immunization with different immunogens, and most immunogens are envisaged to be based on stabilized prefusion HIV Env trimers (*4–7*). This strategy is thought to be necessary both to focus responses to bnAb epitopes and to elicit the SHM needed for bnAb activity.

The advent of the mRNA vaccines against COVID-19 introduced a rapid, effective and promising vaccine platform for human use (*8*). To optimally utilize mRNA for HIV vaccine development, it will be important to evaluate different ways to deliver HIV Env trimers by mRNA. Compared with protein vaccines, mRNA vaccines translate the immunogen protein directly in transfected cells *in vivo*, which enables new modes of antigen delivery, such as membrane-bound antigen delivery, but also increases the burden on antigen design to develop antigens that fold and assemble properly without the benefit of purification. Purified soluble protein trimers employed in many HIV vaccine studies expose an extensive non-glycosylated, non-neutralizing, and typically highly immunogenic surface at the trimer base that is normally occluded on virion-associated, membrane-bound HIV Env (*9*). mRNA delivery of membrane-bound Env trimer has the potential to focus responses to non-base epitopes required for bnAb development. mRNA delivery of membrane-bound HIV Env trimers engineered as germline-targeting priming or first-boosting immunogens has proven effective in mouse models (*10–12*). mRNA delivery of native-like HIV Env trimers has been shown to induce autologous nAb responses in rabbits (*13*) and non-human primates (NHPs) (*14, 15*). Here, we first describe the design and rabbit immunogenicity evaluation of mRNA-delivered soluble and membrane-bound HIV Env native-like BG505-isolate-derived MD39.3 trimers that led to their clinical evaluation in HVTN302 (NCT05217641). We then evaluate NHP B cell, T cell, and antibody responses to mRNA-delivered soluble and membrane-bound MD39.3 trimers compared to a matched MD39.3 soluble protein trimer control with a potent adjuvant.

## Results

### MD39.3 mRNA immunogens were developed and characterized in vitro

For mRNA delivery of BG505 MD39 trimers, we designed soluble gp140 and membrane-bound gp151 forms that contained the (furin cleavage-independent linker) link14 between gp120 and gp41 (*16*), which we refer to as MD39.2, or contained the link14 linker and “congly” mutations to restore the glycosylation sites at positions 241 and 289 (*17*), which we refer to as MD39.3 (**Fig. 1A**). The gp140 versions were truncated at residue 664, as was originally done for BG505 SOSIP (*18*) and later for BG505 MD39 (*19*). For the gp140 constructs, the antigenic profiles, expression yields, thermal stabilities, predominance of native-like conformations as assessed by nsEM, and glycosylation profiles were similar to those for the original cleaved BG505 MD39 gp140 trimer (*19*) (**Fig. S1**). BG505 MD39 and BG505 MD39.3 bind to human CD4 but do not undergo the conformational change necessary to expose the CD4-induced epitope of 17b (**Figs. S1E and S1F**). Additionally, a variant of BG505 MD39.3 containing the G473T mutation to prevent the trimer from binding to human CD4 (CD4KO) (*17*), which could potentially affect its biodistribution was produced and characterized (**Fig. S1F**). The antigenic profiles for the gp151 versions showed appreciable binding to several bnAbs and reduced or undetectable binding to non-nAbs, similar to a cell-surface MD39 gp160 positive control and in contrast to a cell-surface negative control gp120 trimer (**Fig. S2, A to B**).

**Fig. 1.**
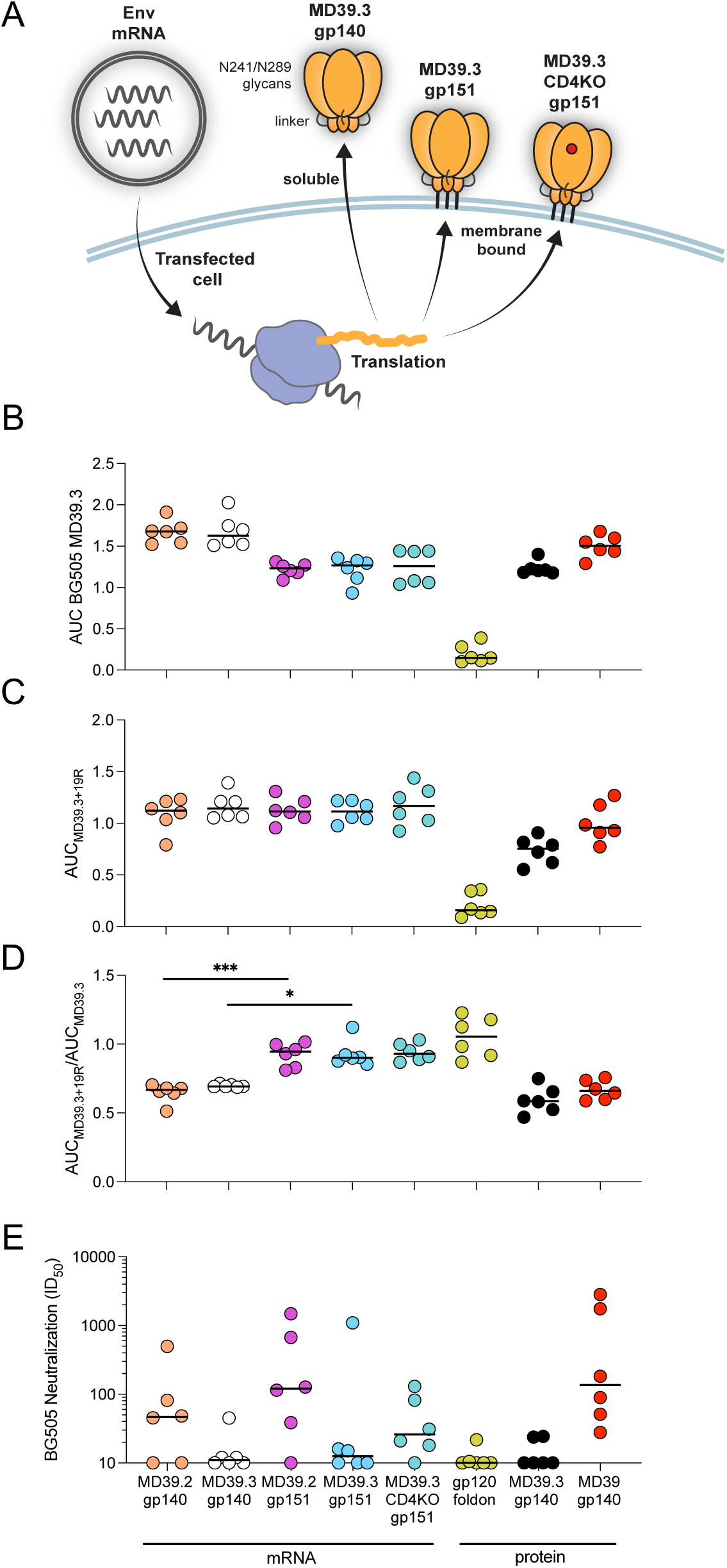
Rabbit serum antibody responses. (**A**) Illustration of MD39.3 mRNA immunogen expression. The cartoon illustrates the in vivo translation of MD39.3 mRNA in transfected cells, producing either soluble or membrane-bound MD39.3 proteins. All MD39.3 variants are designed with furin cleavage independence provided by a Link14 linker and a filled glycan hole achieved through N241/N289 glycans. (**B**) AUC ELISA response for serum antibodies binding to soluble BG505 MD39.3 gp140. (**C**) AUC ELISA response for serum antibodies binding to soluble BG505 MD39.3 gp140 in the presence of the base binding mAb RM19R. (**D**) Ratio of AUC for BG505 MD39.3 gp140 + RM19R over AUC for BG505 MD39.3 gp140 alone. Lower values indicate the presence of more base binding antibodies in the sera. (**E**) Serum neutralization against BG505 T332N pseudovirus. Bars indicate median values for AUC measurements and neutralization data. Each point indicates a single animal (n=6/group). Statistical significance was assessed using the Kruskal-Wallis test, followed by Dunn’s multiple comparisons test. Significance levels were indicated as \**P* < 0.05, \*\**P* < 0.01, \*\*\**P* < 0.001, and \*\*\*\**P* < 0.0001.

### Membrane-bound MD39.3 elicited neutralizing and reduced base antibodies in rabbits

For initial immunogenicity evaluation, we carried out a rabbit study (**Table S1**) in which animals were immunized intramuscularly in the right hind thigh with either 100 µg mRNA-LNPs or 30 µg protein plus 375 µg SMNP adjuvant (*20*) at weeks 0, 8, and 24. All vaccines were based on the BG505 isolate. We tested mRNA-encoded MD39.2 and MD39.3 as both gp140 and gp151, plus the CD4KO variant of MD39.3 gp151. Adjuvanted protein vaccine controls included MD39.3 gp140, fully cleaved MD39 gp140 (which lacked glycosylation sites at 241 and 289), and a negative control gp120 foldon trimer.

We assessed serum antibody binding to MD39.3 at week 26 by ELISA. All groups demonstrated detectable binding to MD39.3, with the gp120 foldon exhibiting barely detectable binding in only a subset of animals, and with groups that received membrane-bound trimer immunogens exhibiting lower binding levels than those receiving soluble trimers (**Fig. 1B**). In the presence of the base-binding monoclonal antibody RM19R (*21*), responses to MD39.3 were similar for all mRNA/LNP groups, including those immunized with soluble or membrane-bound trimers (**Fig. 1C**). By computing the ratio of serum IgG binding in the presence of RM19R compared to without RM19R, we found that approximately 90 to 93% of IgG binding AUC to mRNA-delivered membrane-bound trimers was not blocked by RM19R, whereas only 65 to 69% of the response to mRNA-delivered soluble trimers was not blocked by RM19R (*P* = 0.0008 by Kruskal-Wallis for MD39.2 gp140 vs gp151; *P* = 0.0399 by Kruskal-Wallis for MD39.3 gp140 vs gp151; **Fig. 1D**). This indicated that mRNA delivery of membrane-bound trimers substantially refocused responses away from the base.

The impact of filling glycan holes at positions 241 and 289 in BG505 was evident in the autologous neutralization data. Previous studies have demonstrated that rabbits produce significant levels of autologous neutralizing antibodies targeting the 241-glycan hole on BG505 (*22, 23*). Consistent with expectations, groups immunized with MD39.2 constructs containing glycan holes exhibited higher neutralization titers compared to those receiving MD39.3 constructs, where the glycan holes were filled (**Fig. 1E**). The BG505 gp120 foldon control immunogen showed minimal or no detectable neutralization.

Focusing only on immunogens with intact glycan holes at 241 and 289, we noted that both gp140 and gp151 variants of MD39.2 delivered by mRNA induced comparable neutralization responses to the MD39 adjuvanted protein group (*P* > 0.05 by Kruskal-Wallis for both MD39.2 gp140 or gp151 mRNA vs MD39 gp140 protein; **Fig. 1E**). Combined with the finding from ELISA that gp151 elicited reduced based responses compared to gp140 (**Fig. 1D**), these data suggested that for mRNA delivery, membrane-bound trimers might be superior to soluble trimers.

### EMPEM revealed base-directed and trimer-degrading responses in rabbits

We performed electron microscopy polyclonal epitope mapping (EMPEM) (*24*) as a method to identify which epitopes on MD39.3 are recognized and targeted by serum antibodies in rabbits immunized with different formulations. The immunization groups included MD39.3 gp140 mRNA/LNPs, MD39.3 gp151 mRNA/LNPs, and MD39.3 gp140 protein adjuvanted with SMNP. Analysis of week 10 sera revealed that antibodies from all three groups targeted the base of the trimer (**Fig. 2**). Additionally, all groups exhibited antibodies that caused degradation of the MD39.3 gp140 soluble antigen. The group immunized with the membrane-bound MD39.3 gp151 via mRNA/LNPs demonstrated the most consistent antigen degradation (**Fig. 2**). HIV Env trimer-degrading serum antibodies have been previously observed in EMPEM studies (*25*). When present in substantial amounts, these antibodies pose a challenge for epitope mapping, as the antigen is primarily degraded into protomers. These protomers become saturated with Fabs, complicating their classification through 2D or 3D imaging approaches. To address the potential complications in the week 26 EMPEM analysis, we used an antigen mixture consisting of 80% MD39.3 gp140 stabilized with internal disulfides (*26*) and 20% unmodified MD39.3 gp140. The internal disulfides stabilized the trimer by forming interprotomer covalent bonds without altering the antigenicity of the trimer. This antigen mixture enabled the simultaneous detection of antibodies targeting epitopes on the intact HIV Env trimer and those responsible for trimer degradation. The week 26 EMPEM analysis demonstrated that all immunization groups generated base-specific antibodies. Furthermore, the two groups immunized with mRNA/LNP formulations produced trimer-degrading antibodies. In contrast, the group immunized with MD39.3 gp140 protein combined with the SMNP adjuvant did not exhibit trimer-degrading antibodies at week 26 in quantities sufficient to be detected by EMPEM using the antigen mixture strategy (**Fig. 2**).

**Fig. 2.**
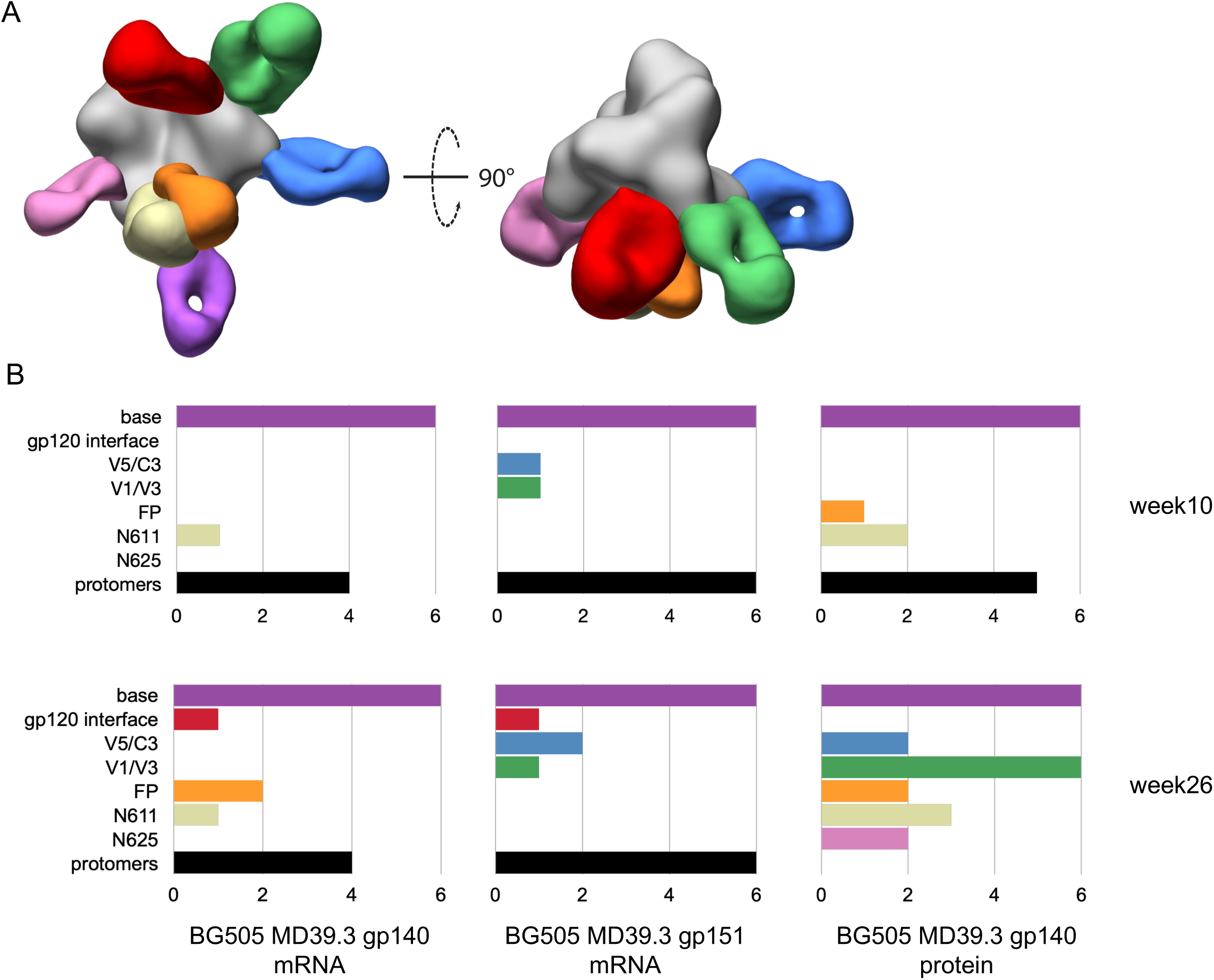
Electron microscopy polyclonal epitope mapping of rabbit serum. (**A**) Composite 3D map representing the epitopes observed in negative stain EMPEM analysis. (**B**) Bar graphs showing the number of animals per with detectable antibodies directed to each specific epitope at 10 and 24 weeks post first immunization. Colored to match (A).

### mRNA-delivered membrane-bound trimers elicited neutralizing antibodies in NHPs

Following demonstration of immunogenicity in rabbits, the MD39.3 immunogens were carried forward to be tested in NHPs. Three immunizations were administered via intramuscular injection bilaterally into the deltoid muscle at weeks 0, 8 and 24 (**Fig. 3A**). Blood and fine needle aspiration of LNs (LN FNA) were collected throughout the study at different time points. A total of 30 NHPs were immunized, organized into 5 groups with 6 animals per group. Three different immunogen designs were delivered as mRNA/LNP: soluble MD39.3 gp140 (Group 1; 100 µg), membrane-bound MD39.3 gp151 (Group 2 & 4; 100 µg & 300 µg), and membrane-bound MD39.3 CD4KO gp151 (Group 3; 100 µg). While the HIV envelope trimer does not have measurable affinity for rhesus macaque CD4, the CD40KO construct was included in the study because it was intended for human trials. Group 5 animals received MD39.3 gp140 protein (100 µg) with SMNP adjuvant (750 µg) as a control.

**Fig. 3.**
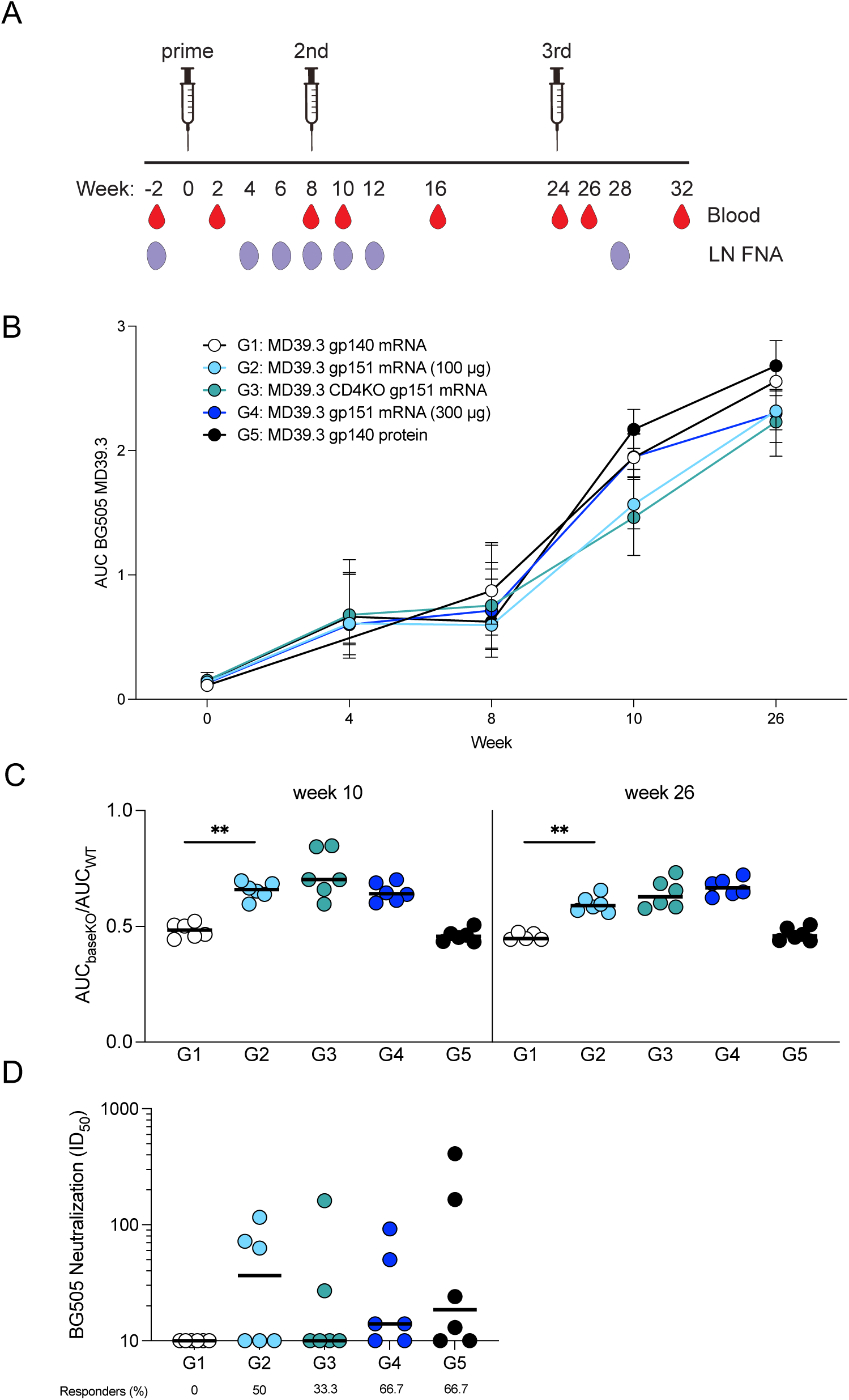
NHP serum antibody responses. (**A**) Three doses of BG505 MD39.3 immunogens were administered at weeks 0, 8 and 24. All doses were administered bilaterally and intramuscularly into the deltoid muscle. PBMCs were isolated from whole blood and LN FNA from axillary LNs. Five groups of six animals per group were immunized with BG505 MD39.3 immunogens. Groups 1-4 received BG505 MD39.3 mRNA immunogens while G5 received BG505 MD39.3 protein plus SMNP adjuvant. The mRNA groups included: soluble MD39.3 (G1), membrane-bound MD39.3 (G2 & G4) and membrane-bound MD39.3 CD4KO (G3). (**B**) Longitudinal AUC ELISA responses for serum antibodies binding to soluble BG505 MD39.3 gp140. (**C**) Ratio of AUC for BG505 MD39.3 gp140 BaseKO over AUC for BG505 MD39.3 gp140 WT. Lower values indicate the presence of more base binding antibodies in the sera. (**D**) Week 26 serum neutralization against BG505 T332N pseudovirus. Bars indicate median values for AUC measurements and neutralization data. Each point indicates a single animal (n=6/group). Groups 1 and 2 were compared for statistical significance using the Mann-Whitney test. Significance levels were indicated as \**P* < 0.05, \*\**P* < 0.01, \*\*\**P* < 0.001, and \*\*\*\**P* < 0.0001.

Serum antibodies binding to MD39.3 gp140 were elicited in all groups of NHPs following the first immunization, with levels increasing over time and after subsequent immunizations (**Fig. 3B**). To differentiate non-base responses from base responses, we designed a BaseKO probe derived from MD39.3, which was used as a capture antigen alongside regular MD39.3. At weeks 10 and 26, a higher proportion of serum antibodies were directed toward the base of the trimer in groups receiving the soluble trimer (G1 and G5) compared with the groups receiving membrane-bound MD39.3 (**Fig. 3C**).

No autologous neutralization against the BG505 pseudovirus was observed in the week 26 sera of animals immunized with MD39.3 gp140 via mRNA/LNPs. In contrast, all other groups demonstrated detectable autologous neutralization activity at week 26 (**Fig. 3D**). Eliminating one of the most commonly targeted neutralizing antibody sites on BG505 Env trimer by filling glycan holes at positions 241 and 289 appeared to substantially reduce the development of neutralizing antibodies. This was most evident by comparing Group 5 to a historical control group that received 100 µg MD39 gp140 protein (with glycan holes at positions 241 and 289) and 750 µg SMNP via subcutaneous bilateral immunizations above the deltoid (50 µg and 375 µg per limb) at 0, 10, and 24 weeks (*20*). All animals immunized with glycan hole-containing MD39 protein developed autologous neutralizing antibodies at week 26, with median titers ∼19-fold higher than after glycan-hole-filled MD39.3 protein immunization (median 353 vs. 19; **Fig. S2C**). In sum, membrane-bound MD39.3 mRNA induced detectable neutralizing responses in some animals compared to soluble MD39.3 mRNA, with responding animals achieving titers comparable to those observed following MD39.3 protein immunization aided by a potent adjuvant.

### mRNA-delivered membrane-bound trimer elicited diverse antibody specificities in NHPs

The same antigen mixture strategy used in the rabbit EMPEM analysis was also employed for the week 26 NHP EMPEM analysis testing soluble MD39.3 gp140 mRNA (G1), membrane-bound MD39.3 gp151 mRNA (G2) and MD39.3 gp140 protein (G5). All three groups analyzed by EMPEM demonstrated base-specific antibodies. However, only G2, which received membrane-bound MD39.3 gp151 via mRNA/LNPs, exhibited detectable antibodies that caused trimer degradation (**Fig. 4**). Additionally, non-base/non-degrading antibodies were identified in both G2 and G5 (**Fig. 4**), which likely contributed to the observed autologous neutralization in these groups (**Fig. 3D**).

**Fig. 4.**
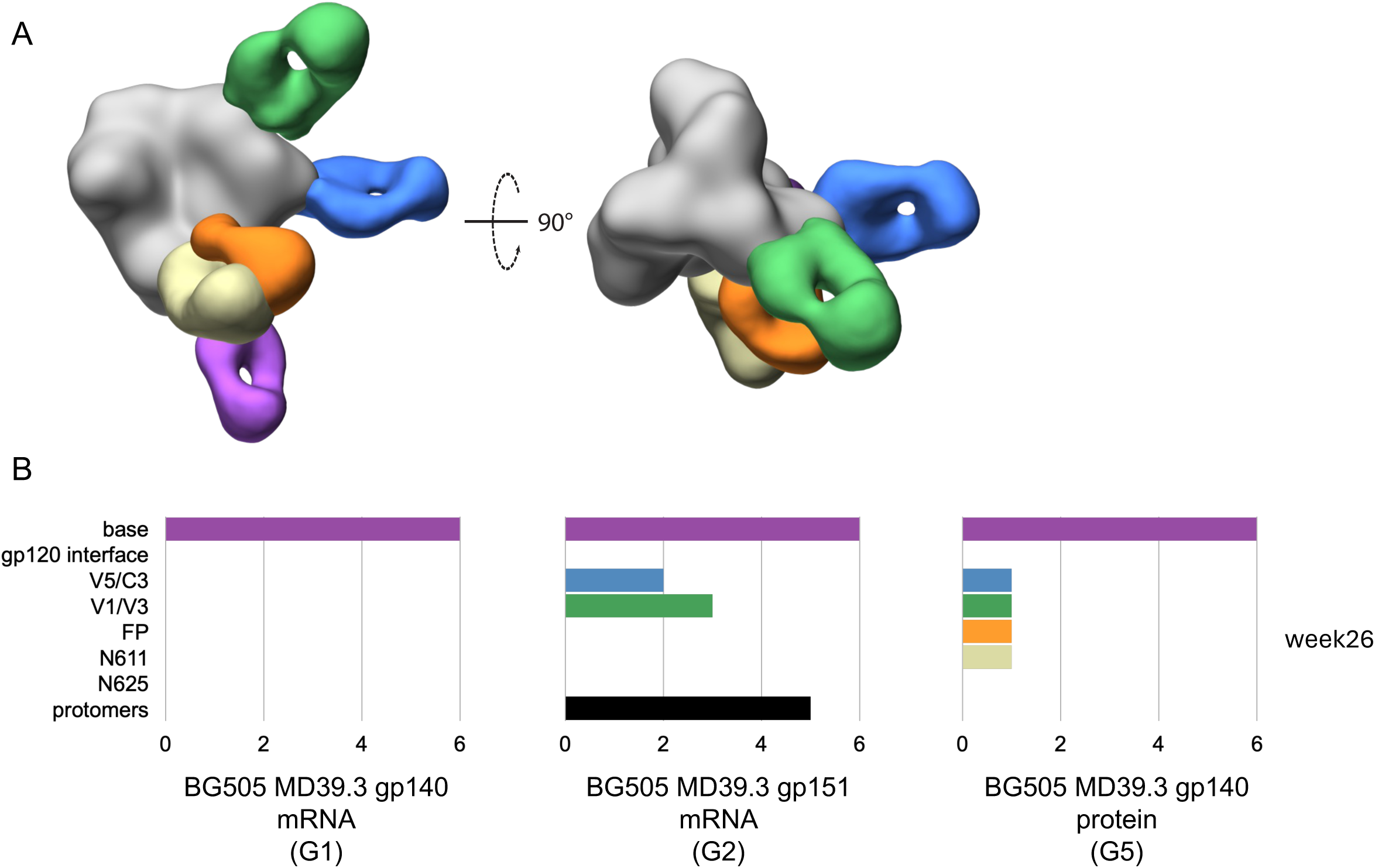
Electron microscopy polyclonal epitope mapping of NHP serum. (**A**) Composite 3D map representing the epitopes observed in negative stain EMPEM analysis. (**B**) Bar graphs showing the number of animals per group with detectable antibodies against each specific epitope at 24 weeks post first immunization. Colored to match (A).

### MD39.3 mRNA induced B cell memory responses in NHPs

For evaluation of NHP B cell responses, we first assessed the immune response in axillary LN FNAs to detect total and Env-binding B cell responses (**Fig. S3, A to C)**. Env-binding germinal center B cells (B_GC_) were detected in some protein immunized animals after 2-3 immunizations, but not in the mRNA groups. The total B_GC_ cell frequencies reflected a similar mild to undetectable response typically seen in a non-draining LN, suggesting that the draining LNs were not successfully accessed by the axillary LN FNAs.

We then investigated the memory B cell (B_Mem_) responses in circulation (PBMCs). Env-binding B_Mem_ cells were detectable post-prime and increased with each dose (**Fig. 5, A to B and fig. S3D**). The fold change of Env-binding B_Mem_ cells over baseline surpassed 100-fold for some animals after each boost (**Fig. 5C**). Env-specific B_Mem_ cells peaked two weeks post-boosts and declined over time (**Fig. 5D**). Compared with soluble versions of MD39.3, the membrane-bound MD39.3 antigens had the highest frequency of non-base binding B_Mem_ cells (BaseKO^+^) across timepoints among the total Env-binding B_Mem_ cells (**Fig. 5E**). Bone marrow (BM) aspirations were collected from select available groups (G1 & G5) more than 1 year since the last immunization. Env-specific BM plasma cells (B_PC_) were detectable for 3 animals per group for G1 and G5 (**Fig. 5F**). In summary, membrane-bound MD39.3 mRNA vaccines induced robust Env-binding B_Mem_ responses after 2 and 3 doses, with reduced frequencies of Env base-binding B_Mem_ cells compared to soluble Env trimer protein or mRNA immunizations.

**Fig. 5.**
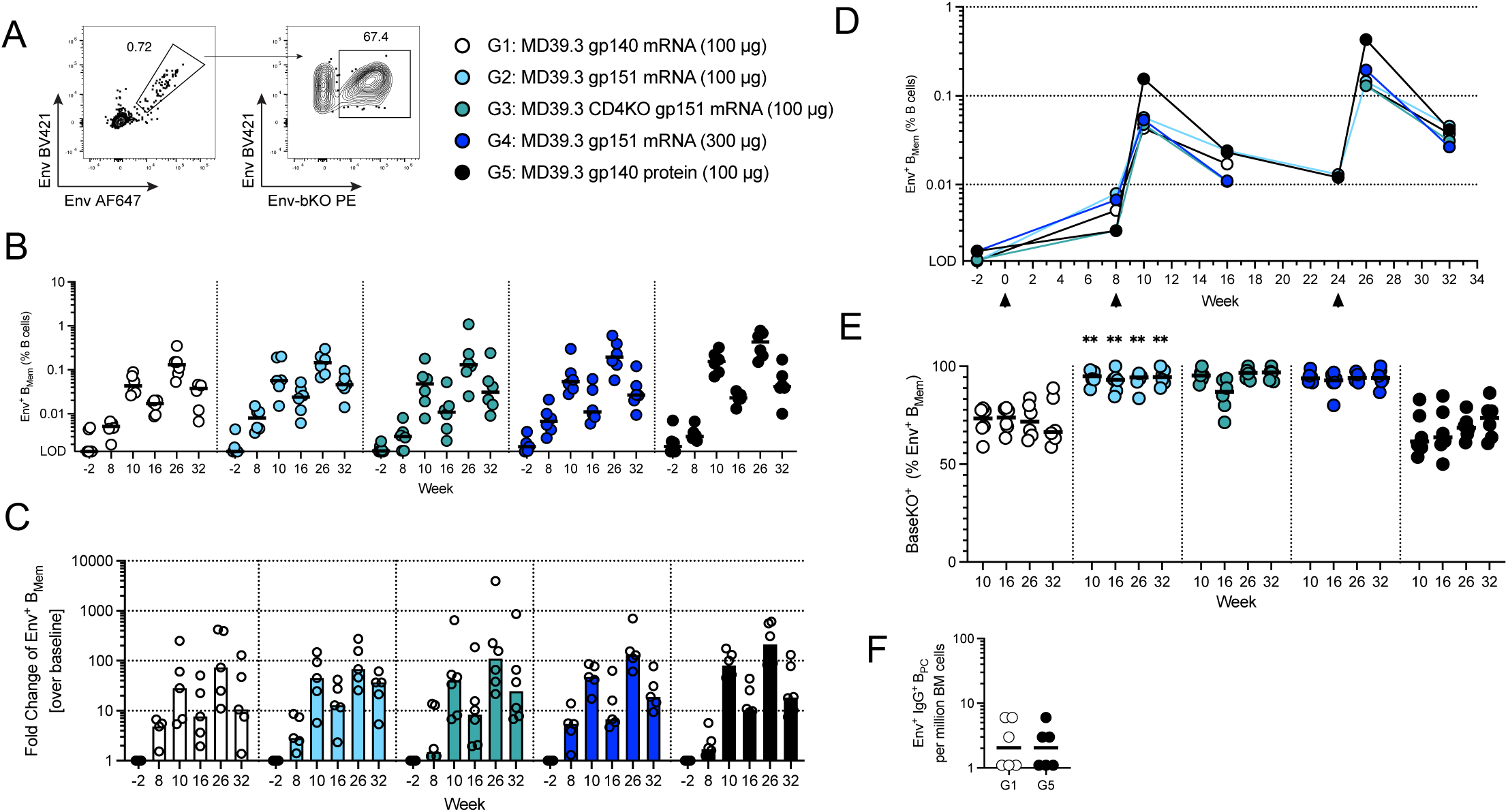
MD39.3-binding B cell responses in NHPs. (**A**) Representative flow plots show CD20^+^IgD^-^IgM^-^Env^+/+^ B_Mem_ cells (left) and BaseKO^+^ within Env^+/+^ (right). (**B**) Frequency of Env^+/+^ B_Mem_ cells in total B cells in PBMCs at different time points are shown. Responses that are lower than the LOD (0.001375) were set at the LOD. (**C**) Fold-change of Env^+/+^ B_Mem_ cells per group and time points over baseline Env^+/+^ B_Mem_ frequencies (pre-immunization week -2). (**D**) Median of Env^+/+^ B_Mem_ cells shown in (B) graphed over time including one additional time point at week 24 (G2 & G5 only). (**E**) Percentage of BaseKO^+^ within total Env^+/+^ B_Mem_ cells in all groups post-boosts. G2 values were compared with corresponding G1 values for statistical significance. (**F**) Amount of Env-specific IgG^+^ B_PC_ in BM aspirates at week 119 (G1 & G5 only). Bars indicate median for B cell frequencies and each point indicates a single animal (n=6/group) except (D) shows median of the respective group. Animals with missing baseline samples were excluded from the fold-change analysis in (C). Groups 1 and 2 were compared for statistical significance using the Mann-Whitney test, followed by the Holm-Šídák multiple comparisons test. Significance levels were indicated as \**P* < 0.05, \*\**P* < 0.01, \*\*\**P* < 0.001, and \*\*\*\**P* < 0.0001.

### MD39.3 mRNA induced antigen-specific T cell responses in NHPs

Based on the B cell and neutralizing responses, we focused our remaining analysis on a single mRNA group: the membrane-bound MD39.3 gp151 mRNA at the 100-µg dose, compared with the adjuvanted MD39.3 protein group. This choice was driven by the superior non-base responses of MD39.3 gp151 compared to the soluble gp140 form, with no obvious benefit at higher doses (300 µg) or from the CD4KO mutation in rhesus macaques. Thus, we tested the capacity of membrane-bound MD39.3 gp151 mRNA (Group 2) and MD39.3 gp140 protein (Group 5) to induce T cell responses post-prime (week 2) and post-boost (week 26) by activation-induced marker (AIM) and intracellular cytokine staining (ICS) assays. Both groups induced similarly robust Env-specific AIM CD4 T cell responses in circulation (**Fig. 6, A to B and fig. S4A**). The combination of AIM plus ICS showed that priming with mRNA-delivered membrane-bound MD39.3 generated Env-specific CD4 T cells capable of producing IFNγ, TNF, granzyme B, or IL-2 (**Fig. S4B)**. Env-specific circulating T follicular helper (cT_FH_) cells were induced by mRNA-delivered membrane-bound MD39.3 priming to levels comparable to protein immunization (**Fig. S4C**). Env-specific CD8 T cell responses were detected in the majority of the mRNA-delivered membrane-bound MD39.3 animals (**Fig. 6, C to D**). Overall, after a single immunization, mRNA-delivered membrane-bound Env generated detectable CD8 T cell responses and consistent Env-specific CD4 T cell responses with diverse functionalities.

**Fig. 6.**
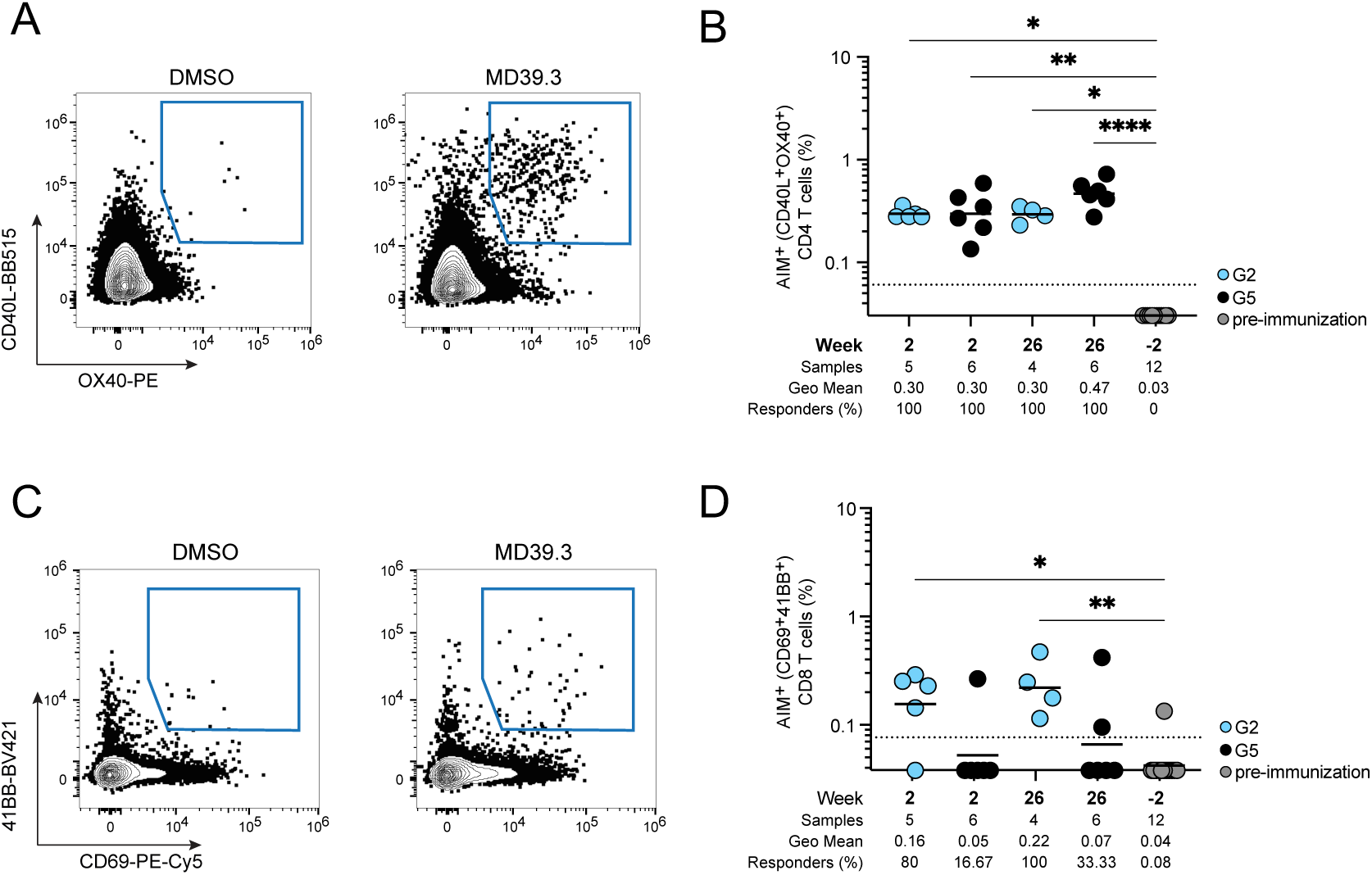
MD39.3-specific T cell responses in NHPs. (**A**) Env-specific AIM^+^ CD4 T cell responses of G2 and G5 at week 2 (post-prime) and week 26 (post-third). (**B**) Env-specific AIM^+^ CD8 T cell responses of G2 and G5 at week 2 (post-prime) and week 26 (post-third). Data are shown as background subtracted. Non-responder samples are set at baseline. The dotted black line indicates the limit of quantification (LOQ). Bars represent geometric mean and each point indicate a single animal (n=6/group). Animals with missing or poor-viability samples were excluded from the analysis. Statistical significance was assessed using the Kruskal-Wallis test, followed by Dunn’s multiple comparisons test. Significance levels were indicated as \**P* < 0.05, \*\**P* < 0.01, \*\*\**P* < 0.001, and \*\*\*\**P* < 0.0001.

### Sequencing of antigen-binding B_Mem_ cells revealed mutational and clonal dynamics

Single B cell receptor (BCR) sequencing was utilized to investigate the sequences and mutational patterns of Env-binding B_Mem_ clones. We sorted and sequenced Env-binding B_Mem_ cells in PBMCs from mRNA-delivered membrane-bound MD39.3 immunized animals (Group 2) and MD39.3 protein immunized animals (Group 5) at five timepoints. Post-prime at week 8, Env-binding B_Mem_ cells exhibited relatively high SHM in both vaccination groups, with over 95% of the recovered Env-binding B_Mem_ cells being mutated (**Fig. 7A**). Two weeks post-boost, (week 10) the frequencies of Env-binding B_Mem_ cells increased substantially (**Fig. 5B, 5D, 7A**), with > 99% exhibiting SHM. Median SHM in Env-binding B_Mem_ cells was greater at week 24 than week 10 for animals in both vaccination groups (*P* < 0.0001 by Mann-Whitney for both groups 2 and 5; **Fig. 7A**). Two weeks after the third dose (week 26), SHM of the Env-binding B_Mem_ cells was similar to week 24. The robust induction Env-binding B_Mem_ cells with SHM confirmed that GCs were induced in these animals, but the draining LN was not sampled in the FNAs. Overall, Env-binding B_Mem_ cells accumulated substantial SHM in their BCR after the prime and boost immunizations, indicating ongoing affinity maturation.

**Fig. 7.**
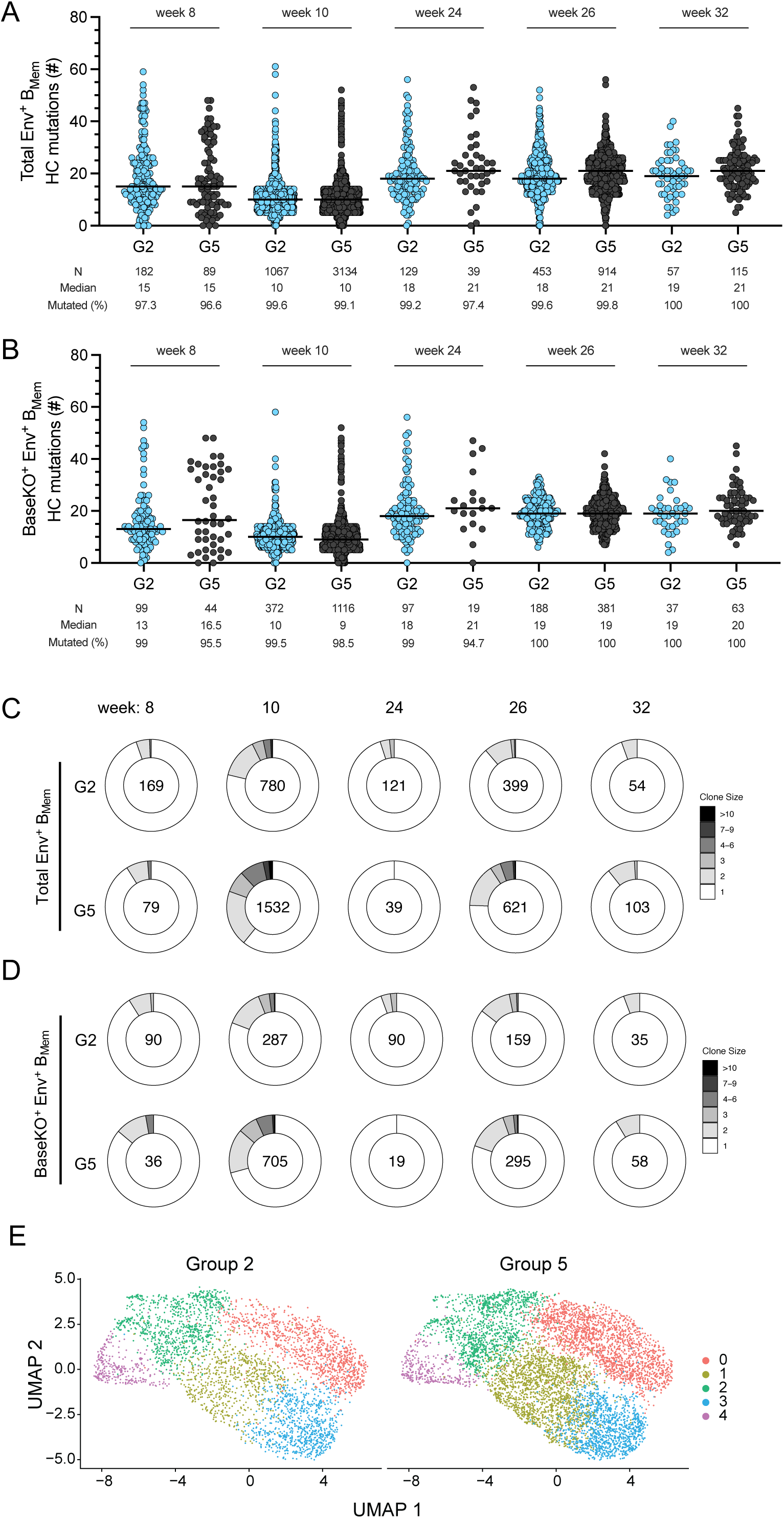
Single cell analysis of MD39.3-binding B_Mem_ cells in NHPs. (**A**) Heavy chain mutations in total Env-binding B_Mem_ cells were assessed for G2 and G5 at week 8, 10, 24, 26 and 32. (**B**) Heavy chain mutations on non-base-Env-binding B_Mem_ cells (BaseKO^+^) for the same groups and timepoints as in (A). (**C**) Total Env-binding B_Mem_ clonal families were analyzed for groups 2 and 5 across timepoints. Numbers within the Donut plot indicate total number of clonal families detected for each measurement. (**D**) Non-base-Env-binding B_Mem_ clonal families were analyzed for the groups and timepoints shown in (C). **(E**) Uniform manifold approximation and projection (UMAP) visualization of single-cell gene expression profiles identifying clusters among Env-binding B_Mem_ cells sorted from PBMCs for G2 and G5. Bars indicate median for mutational analysis and each point indicates a single B cell sequence.

We expressed several monoclonal antibodies (mAbs) from select BCR sequences of Group 2 animals at week 26 and mapped their epitope specificities using SPR and a panel of mutant MD39.3 antigens (**Fig. S5A**). Among the mAbs that bound to MD39.3, three targeted the base region, two targeted the C3/V5 region, and two targeted the V1/V3 region. Other mAbs exhibited high-affinity binding to MD39.3 but could not be mapped to a specific epitope or region. mAbs with an affinity of 100 nM or better for MD39.3 were further assessed for neutralization potential. Only the two mAbs targeting the V1/V3 region demonstrated the ability to neutralize autologous BG505 pseudovirus and several epitope mutants (**Fig. S5B**). The lack of neutralization despite high-affinity binding to MD39.3 observed for many mAbs suggested that these mAbs may be recognizing peptide epitopes present on MD39.3 but absent or shielded on the BG505 pseudovirus (i.e Link14 or the V3 tip). To address this potential, we assessed the ability of the mAbs to bind to a V3 peptide or the Link14 peptide and found that none of the mAbs bound either peptide (**Fig. S5C**). Overall, the immunogen induced mAbs with affinities in the nanomolar range, including some with neutralizing capabilities.

Inclusion of a barcoded BaseKO probe in the B_Mem_ sorts allowed discrimination between Env base-binder and non-base binder B_Mem_ cells. Non-base-binding B_Mem_ cells possessed similar SHM kinetics compared to total Env-binding B_Mem_ cells, with > 95% of non-base-binding B_Mem_ cells mutated post-prime (**Fig. 7B**). The interclonal diversity of Env-binding B_Mem_ cells was high. Larger clonal families were detected at the peak post-boost timepoints at week 10 and 26 (**Fig. 7, C to D**). V gene usage was overall similar between both groups (**Fig. S6A**), as were CDRH3 lengths (**Fig. S6B**). Sorted Env-specific B_Mem_ phenotypic characteristics after mRNA versus protein immunization was assessed by whole transcriptomic gene expression profiling. Five B_Mem_ clusters were identified (**Fig. S7**). All five B_Mem_ clusters were represented after mRNA or protein immunization (**Fig. 7E**), as well as at every time point (**Fig. S7**).

## Discussion

No HIV vaccine has been approved to date, and major efforts are focused on inducing protective immune responses through broadly neutralizing antibodies (bnAbs). However, naïve (germline) precursor B cells capable of maturing into bnAbs are typically rare and rely on competitive recruitment into the GC response. Germline-targeting immunogens represent a promising approach for the initial recruitment of these rare B cell precursors (*4, 16, 27, 28*). Following successful recruitment, the newly differentiated B_GC_ and B_Mem_ cells require sequential immunizations with increasingly native-like immunogens to guide them through rounds of clonal expansion and SHM. This process aims to ultimately refine the bnAb precursors into functional bnAbs. The immunogen MD39.3 tested here could serve as a potential candidate for use as a polishing immunogen in these germline-targeting strategies. The data indicate that MD39.3 mRNA functions as an effective immunogen in preclinical mammalian models, enabling subdominant epitope responses when expressed in its membrane-bound form compared to its soluble form. The comparison between mRNA and protein underscores the high standard set by decades of research dedicated to optimizing protein vaccine design and adjuvant strategies. In this study, immunization with membrane-bound MD39.3 mRNA elicited B_Mem_, neutralizing antibody, and CD4 T cell responses comparable to those achieved with MD39.3 protein formulated with SMNP adjuvant. Additionally, no concerning events were observed in any of the animals. Thus, MD39.3 mRNA offers several advantages over MD39.3 protein, including: 1) eliminating the need for an additional adjuvant, 2) reducing exposure of the trimer base, and 3) simpler and more cost-effective manufacturing. In summary, our findings demonstrate that membrane-bound MD39.3 mRNA represents a promising platform for vaccine development strategies and suggest more generally that for mRNA delivery of HIV trimers at any stage of a germline-targeting vaccine regimen, membrane-bound trimer is likely preferred over secreted soluble trimer.

A comparison here to historical NHP MD39 immunization data indicated that the glycan hole filling of MD39.3 makes a substantially more challenging target for BG505 autologous neutralizing antibody generation, which is a useful reference point when comparing across diverse BG505 Env trimer immunization study results, including in humans.

The robust detection of Env-binding B_GC_ cells in some animals only in the protein immunization group (Group 5) may indicate differences in NHPs in drainage patterns and localized immune responses between vaccine platforms. Previous studies have suggested that intramuscularly injected immunogens exhibit more restrictive LN drainage compared to subcutaneous administration (*29, 30*). Nonetheless, the data herein indirectly demonstrate robust GC responses occurred after mRNA priming and boosting, as evidenced by almost all B_Mem_ exhibiting SHM, with SHM levels comparable to those for protein immunization plus SMNP, an adjuvant that has been shown to induce robust GC responses (*4, 20, 31, 32*). Previous studies have shown that intramuscularly delivered COVID-19 mRNA vaccines induce robust and long-lasting GC responses in humans (*33*). Other recent studies have found evidence that LN drainage for intramuscular mRNA vaccination in NHPs may differ from protein immunization (*34*). Long-lasting GC responses to a priming immunization are preferred for enhancing affinity maturation and promoting competitive engagement of rare B cell clones targeting subdominant epitopes (*31, 35, 36*). Ongoing investigations aim to better understand how mRNA vaccines drain in NHPs and inform on GC responses following immunization.

We observed a strong induction of Env-binding B_Mem_ cells following immunization, with peak responses detected in NHPs two weeks after each boost, followed by a decline in subsequent weeks, consistent with antigen-binding B_Mem_ kinetics in NHPs after two doses of a COVID-19 mRNA vaccine (*37*). This suggests that NHPs may exhibit distinct B_Mem_ kinetics compared to humans, possibly due to a pronounced but short-lived expansion of B_Mem_ cells shortly after boosting, regardless of the vaccine platform.

Both membrane-bound MD39.3 mRNA and protein immunizations induced Env-specific CD4 T cell responses. Interestingly, low levels of Env-specific CD8 T cell responses were also observed, with a slight trend toward higher responses in the mRNA vaccine group compared to the protein group. COVID-19 mRNA vaccines induce CD8 T cell responses in humans (*38–40*). The presence of Env-specific CD8 T cells in mucosal tissues could synergize with nAbs to provide protection against HIV infection (*41*). These responses may also be useful in therapeutic vaccines. Thus, understanding how to induce and enhance antigen-specific CD8 T cell responses elicited by mRNA vaccines remains an important goal for future research.

Limitations: In addition to limitations discussed above, EMPEM provides qualitative insights into various antibody-antigen binding modes but does not provide quantitative data on the proportion of the overall immune response targeting a specific site. As a native-like immunogen, MD39.3 alone is insufficient to elicit broad neutralization.

## Materials and Methods

### Study design

The primary objectives of this study were to investigate the preclinical immunogenicity of BG505 MD39.3 HIV envelope trimer delivery using mRNA/LNP in mammalian models. Rabbits were used as a small mammalian animal model, while NHPs served as a physiologically relevant model for humans to assess translational potential. The MD39.3 mRNA immunogens were tested in two formats: soluble and membrane-bound. All animal procedures complied with NIH guidelines and were approved by the respective Institutional Animal Care and Use Committees (IACUCs). Group sizes were predetermined to include six animals per group, based on prior power analysis (*42*), to ensure sufficient statistical power for reliable measurement of immunological responses and detection of meaningful differences in a single experiment. The population from which the animals were selected was defined based on availability and suitable age groups for the study. Animals were randomly assigned to experimental groups while ensuring balanced sex representation. The study was not blinded. Data collection continued until all planned endpoints were reached. The animals were monitored daily by clinical veterinarians, and physical examinations were conducted during each anesthetic access. No concerning events were reported in the animals. All animals, including outliers, were included in the analysis, with the exception of animals with poor viability PBMCs in the T cell analysis (Fig. 6 and fig. S4). Primary endpoints included safety and immunogenicity measures such as antibody titers, neutralization responses, and antigen-specific B and T cell responses.

### Protein Production

BG505 MD39.2 gp140 and BG505 MD39.3 gp140 trimer immunogens; His-tagged trimer ELISA, SPR, and EMPEM reagents; and His-Avi-tagged biotinylated trimer probes were produced by transient co-transfection of HEK-293F cells (Thermo Fisher) as previously described (*19*). For BG505 MD39 gp140, co-transfection using a plasmid encoding human Furin was performed to ensure gp120/gp41 cleavage as previously described (*19*). His-tagged and His-Avi-tagged trimers were purified by immobilized metal ion affinity chromatography (IMAC) using HisTrap excel columns (Cytiva) followed by size-exclusion chromatography (SEC) using a Superdex 200 Increase 10/300 GL column (Cytiva). His-Avi-tagged trimers were biotinylated using BirA (Avidity) according to the manufacturer’s instructions and purified again to remove excess biotin using SEC. BG505 MD39 gp140, BG505 MD39.2 gp140, and BG505 MD39.3 gp140 trimers were purified by 2G12 affinity chromatography followed by SEC. Immunogen preps were confirmed to contain < 5 EU/mg of endotoxin using an Endosafe instrument (Charles River). Monoclonal IgGs were produced in house as previously described (*43*). Monoclonal Fabs produced and purified as previously described (*4*). The following mutant versions of BG505 MD39.3 were produced for use in ELISA, EMPEM, SPR, and flow cytometry:

- BG505 MD39.3 CC5 = BG505 MD39.3 + E49C-L555C and A73C-P561C
- BG505-MD39.3 V1/V3-KO = BG505 MD39.3 +133aN and 136aA
- BG505-MD39.3 C3/V5-KO = BG505 MD39.3 + T461N, S463T, T464N, and E466S
- BG505-MD39.3 BaseKO (sorting probe) = BG505 MD39.3 + R500A, Q658K, and A662E
- BG505-MD39.3 BaseKO (SPR/ELISA) = BG505 MD39.3 + R500A, C605T, Q658T, L660N, A662T, L663C, D664N, 665G, and 666T

### Cell Surface Antigenicity

Cell surface antigenicity of membrane-bound HIV Env trimers was assessed using flow cytometry. 25 μg of plasmid DNA encoding membrane-bound HIV Env trimers in 833 μL of Opti-MEM media (Thermo Fisher) was added to 833 μL of Opti-MEM media (Thermo Fisher) containing 50 μL of 293Fectin (Thermo Fisher), mixed and incubated at RT for 15 to 20 minutes. Transfection reactions were added to 25 mL of HEK293F cells at 1 million cells/mL in FreeStyle 293F media. All transfections were carried out in duplicate. Transfected cells were incubated at 37°C and 8% CO_2_ on an orbital shaker platform at 125 rpm for 2 days. Transfected cells were transferred to deep-well 96-well plates (1mL per well) and spun down at 650 x g for 5 minutes. Cells were resuspended in 110 μL FACS buffer (2% FBS in PBS) containing 10 μg/mL primary Fabs (PGT145, PGT151, VRC01, PGT121, PGT128, 10E8, F105, B6, 19b, or PBS control), transferred to U-bottom 96-well plates, and incubated on a plate shaker for 1 hour at 4°C. Cells were washed twice with FACS buffer and resuspend in 50 μL FACS buffer containing 0.05 μL SYTOX Green Dead Cell Stain (ThermoFisher) and 0.25 μL AF647 AffiniPure Goat Anti-Human IgG F(ab’)₂ fragment specific (Jackson ImmunoResearch, Catalog #109-605-006). Stained cells were incubated for 20 minutes at RT in the dark and washed twice with FACS buffer. Cells were resuspended in 150 μL FACS buffer. Cells were analyzed on a NovoCyte 3000 flow cytometry (Agilent) using BD FACS Diva 6 software (BD Biosciences). Approximately 50k live cells were acquired per well. Data were analyzed using FlowJo v10.8 and plotted using Prism (v10.4.1; GraphPad).

### Biolayer interferometry (BLI)

Soluble gp140 immunogen antigenicity was assessed using BLI as previously described (*44*) using a panel of bnAbs and non-neutralizing mAbs. A soluble BG505 gp120 foldon trimer was used as a control to show the antigenic profile of a non-native trimer (low binding to quaternary specific bnAbs and high binding to non-neutralizing mAbs). The impact of binding human CD4 on the antigenicity of the soluble gp140 immunogens was assessed using BLI. Biotinylated versions of BG505 SOSIP, BG505 MD39, BG505 MD39.3, and BG505 MD39.3-CD4KO were captured on streptavidin biosensors (Satorius) at 25 µg/mL in kinetics buffer (1X PBS, 0.01% (w/v) BSA, 0.002% (v/v) Tween-20) until a response shift of 0.5 nm was reached. After acquiring a baseline in kinetics buffer alone, biosensors were transferred to wells containing 2 µM human soluble CD4 (sCD4) (*45*) or kinetics buffer alone and allowed to saturate for 2 minutes. The biosensors were transferred to wells containing 2 µM 17b IgG or kinetics buffer alone for additional 2 minutes to assess binding of 17b IgG after CD4 induced conformation change. For the peptide BLI experiment, biotinylated versions of the Link14 (VGSHSGSGGSGSGGHAAAGGAGK-biotin) and V3 (TRPNNNTVKSIRIGPGQAFYYTGGAGK-biotin) peptides were produced by GenScript and loaded onto streptavidin biosensors (Satorius) at 10 µg/mL in kinetics buffer. After acquiring a baseline in kinetics buffer alone, biosensors were transferred to wells containing 1 µM various IgGs in kinetics buffer. The V3 specific mAb 19b was included as a positive control along with negative control bnAbs BG18 and PGT145.

### Negative stain electron microscopy

MD39.3 SOSIP was diluted in Tris-buffered saline to 0.02 mg/mL and adsorbed onto a carbon-coated and plasma cleaned copper mesh EM grid. Following staining for 45 s with 2% (w/v) uranyl formate, approximately 100 micrographs were collected on a Thermo Fisher Talos F200C equipped with a Ceta 16M camera using EPU software. Particles were picked, extracted and subjected to 2D classification using Relion 4.0 (*46*). The number of native-like trimers was calculated by summing particles belonging to 2D class with clear, compact trimers and dividing by the total number of particles in the 2D classification job. For BG505 or MD39.3 complexes with human sCD4, each trimer was incubated with 6X molar excess of sCD4 to trimer for 3 hr at RT. Grid preparation, data collection and analysis steps were identical to those above for MD39.3 alone.

### Glycan Occupancy

Site specific glycan profiling was conducted as previously described (*47*). The degree of glycan occupancy and proportion of glycans that were complex and oligomannose/hybrid type were determined.

### mRNA immunogens

Amino acid sequences encoding BG505 MD39.2 and BG505 MD39.3 immunogens (gp140 and gp151) were provided to Moderna for production as mRNA/LNP immunogens.

### Animals and immunizations

Female New Zealand white rabbits (ages at start of study: range, 2.8 to 33 months; average age, 6.0 months; weights at start of study: range, 1.6 to 4.1 kg; average weight, 2.7 kg) were housed and immunized at Pocono Rabbit Farm (Canadensis, PA). Animals were immunized intramuscularly with a single 500 µl injection in the right thigh. Doses per injection were: 30 µg (protein), 375 µg (SMNP adjuvant), 100 µg (mRNA/LNP), all diluted in PBS. This animal experiment was conducted in compliance with the Animal Welfare Act Regulations (9 CFR), U.S. Public Health Service Office of Laboratory Animal Welfare (OLAW) Policy on Humane Care and Use of Laboratory Animals, and AAALAC accreditation. The immunization protocol (20G131) was reviewed and approved by the Animal Care and Use Committee of the Pocono Rabbit Farm and Laboratory.

Non-human primate study animals were 3-4 years old male and female with a weight range of 5.2-7.4 kg Indian-origin rhesus macaques (*Macaca mulatta*). Animals were bred and maintained at Emory National Primate Center at Emory University in strict accordance under protocols 202000087 and 2021000197 approved by the Animal Care and Use Committee (IACUC) at Emory. Animal care facilities are accredited by the U.S. Department of Agriculture (USDA) and the Association for Assessment and Accreditation of Laboratory Animal Care (AALAC) International. Animals were tested negative for simian immunodeficiency virus. All immunizations were administered intramuscularly (IM) into the left and right deltoid muscle. The total dose for all MD39.3 mRNA vaccines was 100 µg (except for Group 4 the dose was 300 µg). The total dose for MD39.3 protein was 100 µg plus 750 µg SMNP adjuvant. Blood and LN FNAs were collected at indicated time points from the animals. PBMCs and plasma were separated from whole blood and serum was collected independently. BM aspirates were collected only from available animals (Group 1 and 5). Animals were treated with anesthesia (ketamine 5-10 mg/kg or telazol 3-6 mg/kg) and analgesics for IM immunizations, LN FNA, BM aspirates, and blood draws as per veterinarian recommendations and IACUC approved protocols. After completion of the proposed study and approval of the vet staff, animals were released to the center for reuse by other researchers.

### ELISA

On Day 1, human PGT128 capture antibody was coated onto ELISA plates (Corning™ 96-Well Half-Area Plates, Catalog #3690) at a concentration of 1 or 2 μg/mL in 25 μL PBS (pH 7.4; Thermo Scientific, Catalog #10010-023) per well. Plates were incubated overnight at 4°C. On Day 2, plates were washed three times (3X) with PBST (PBS + 0.2% Tween-20) and blocked for 1 hour at room temperature (RT) with PBST containing 5% skim milk (BD Difco™ Skim Milk, Catalog #232100) and 1% Fetal Bovine Serum (FBS; Thermo Fisher, Catalog #16000044). After blocking, plates were washed 3X with PBST, and 25 μL of ELISA antigen (0.3, 1, or 2 μg/mL) was added per well. Plates were incubated for 2 hours at RT, followed by 3X washes with PBST. To assess trimer bottom-directed non-specific responses, a competition ELISA was performed using trimer base-targeting RM19R antibody (*21*). Specifically, 25 μL of RM19R IgG at 20 μg/mL was added per well, and plates were incubated for 1 hour at RT. Subsequently, serum serially diluted (starting at 1:100, 3-fold dilutions) in blocking buffer (PBST + 1% FBS) was added and incubated for 1 hour at 37°C with 80% humidity. Plates were washed 3X, and 25 μL of secondary antibody was added as follows: for rabbit serum, Peroxidase AffiniPure Donkey Anti-Rabbit IgG (H+L) (Jackson ImmunoResearch, Catalog #711-035-152), diluted 1:5,000 in PBST + 1% FBS was used; for NHP serum, Peroxidase AffiniPure™ Donkey Anti-Human IgG, Fcγ fragment specific (Jackson ImmunoResearch, Catalog #709-035-098), diluted 1:5,000 in PBST + 1% FBS was used. Plates were incubated for 1 hour at RT, washed 3X, and developed using TMB Chromogen Solution (Thermo Fisher, Catalog #002023) diluted 1:4. After 6 minutes, the reaction was stopped by adding 25 μL of 0.5 M H₂SO₄. Absorbance was measured at 450 nm and 570 nm using a BioTek Epoch 2 Microplate Reader (BioTek). Background subtraction was performed by subtracting the 570 nm value from the corresponding 450 nm value. The resulting data were analyzed using Prism software (v10.4.1; GraphPad) to calculate the Area Under the Curve (AUC). AUC was determined using the trapezoidal method, which connects adjacent points with straight lines and sums the areas under these segments to approximate the total curve area.

### Neutralization assay

Pseudovirus neutralization assays were performed as previously described using the BG505 T332N pseudovirus (*48*).

### Electron microscopy-based polyclonal epitope mapping (EMPEM)

Polyclonal Fabs were prepared as previously described (*49*). Briefly, IgG was isolated from heat-inactivated serum samples using a HiTrap MAbSelect PrismA protein A column (Cytiva). Purified polyclonal IgG was digested using papain to generate polyclonal Fab, and Fc and undigested IgG was removed by incubating with CaptureSelect IgG-Fc multispecies affinity resin (Thermo Fisher). 0.5 mg of each polyclonal Fab sample was complexed with 15 µg of either BG505 MD39.3 or 15 µg of an 80/20 mixture of BG505 MD39.3 CC5 (containing 2 engineered disulfides: 49C-555C, 73C-561C) and BG505 MD39.3. Samples were incubated overnight at 20°C. Complexes were purified over a Superdex 200 Increase size exclusion column (Cytiva). EM grids were prepared by diluting complexes were to ∼0.03 mg/mL (based on trimer concentration) in 1X TBS. A 3 µL drop of sample was applied to a glow-discharged carbon-coated copper mesh grid for 10 s, blotted with Whatman #1 filter paper, and a 3 µL drop of 2% (w/v) uranyl formate solution was applied for 45-60 s, followed by blotting. Grids were imaged using an FEI Tecnai Spirit microscope operating at 120 keV, equipped with a TVIPS TemCam F416 camera, and 52,000x magnification (resulting in a 2.06 Å pixel size). Data acquisition was automated using Leginon (*50*) and processed using Relion 4.0 (*46*). Visualization and image generation was performed using UCSF ChimeraX (*51*). Representative EM maps have been deposited to the Electron Microscopy Data Bank (EMDB).

### Flow cytometry

Cryopreserved PBMCs or LN FNAs were thawed and washed in R10 media. Tetramerized fluorescent probes for MD39.3 and MD39.3 CD4KO and corresponding BaseKO were prepared by mixing each biotinylated probe with streptavidin-conjugated fluorophores in stepwise addition and incubated in the dark at room temperature for a total of 45 minutes. MD39.3 BaseKO tetramers were added first to the cells and incubated for 20 minutes on ice, followed by addition of MD39.3 tetramers for 30 minutes and lastly adding the complete surface Ab staining cocktail for 30 minutes on ice. Stained cells were washed and analyzed on the Aurora (Cytek Biosciences) or sorted on the FACSymphony S6 (BD Biosciences). Sorted week 24 samples (group 2 & 5) analysis was included with other sample data analyzed on the Aurora included in Fig. 2D for longitudinal responses. Antibody panels are summarized in Table S2 to S4. For single cell sequencing analysis, TotalSeq-C anti-human Hashtag antibodies were used to multiplex samples. TotalSeq-C PE Streptavidin and custom TotalSeq-C BV421 Streptavidin conjugates were used for probe labeling (BioLegend). The limit of detection (LOD) for Env-binding B_GC_ cells was calculated based on the median of (3/(number of B cells collected)) from the LN FNA samples at the pre-immunization timepoint and for Env-binding B_Mem_ cells was calculated based on the median from the PBMC samples at the pre-immunization timepoint.

### Antigen-specific T cell AIM/ICS assay

Cryopreserved PBMCs were thawed and washed in R10 media. Cells were counted and seeded at 1 x 10^6^ per well in a round-bottom 96-well plate in presence of 5 µg/mL MD39.3 peptide pool, 1 ng/mL staphylococcal enterotoxin B (SEB) used as a positive control and DMSO as a negative control (an equimolar amount of DMSO is present in the peptide pool, which was plated in duplicate). Prior to addition of peptides, cells were blocked with 0.5 μg/mL anti-CD40 mAb (Miltenyi Biotec) and incubated with anti-CXCR5 and CCR7 15 min at 37° C. After 24 hours of incubation, intracellular transport inhibitors (0.25 μL/well of GolgiStop and GolgiPlug (BD Biosciences)) and the AIM marker antibodies were added to the samples and incubated for an additional 4 hours. At the end of the culture, cells were washed and stained with surface markers for 30 minutes at 4°C before fixation and permeabilization and subsequent intracellular cytokine staining at RT for 30 min. Stained cells were washed and analyzed on the Aurora (Cytek Biosciences). Antibody panels are summarized in Table S5. Antigen-specific CD4 and CD8 T cells were calculated by subtracting the background data (average DMSO response). A minimum threshold for DMSO levels was set at 0.005%. For each sample, the stimulation index (SI) is calculated as the ratio of the frequency of AIM+ cells in the MD39.3 stimulated condition compared to the averaged DMSO response for the same sample. The limit of quantification (LOQ) was set at the geometric mean of all DMSO samples. Samples with an SI < 2 for CD4 T cell or 3 for CD8 T cell responses and/or with a background-subtracted response <LOQ were considered non-responders. Non-responder samples were set at the baseline level.

### B cell receptor sequencing and analysis

Sorted Env-binding B_Mem_ cells, Single Cell VDJ 5’ Gel Beads and partitioning oil were loaded into the designated wells of the Chromium Next GEM Chip K and run on the Chromium Controller for the generation of single cell droplets. GEX, VDJ and Feature Barcode libraries were generated using the Chromium Next GEM Single Cell 5’ Reagent Kits v2 or v3 (10X Genomics) according to manufacturer’s recommendations. Custom primers specific for NHP immunoglobulin constant regions were designed for the VDJ amplification step. Samples were sequenced on the NovaSeq 6000 (Illumina). Raw sequences were assembled using the de novo option in CellRanger (10X Genomics). Assembled contigs were annotated using IgBLAST on a custom rhesus macaque reference and further processing was done using the Change-O pipeline from the Immcantation portal (*52*). The SHazaM package was used to determine mutations in the BCR sequences and clonal related sequences were clustered with DefineClones.py using the appropriate threshold. Sequences with IgM or NA isotypes were excluded from the SHM analysis.

### Surface Plasmon Resonance (SPR)

We measured kinetics and affinity of antibody-antigen interactions on a Carterra LSA using a low-capture IgG method as previously described (*43*). Epitope mapping was performed using the knockout versions of BG505 MD39.3.

### Single cell transcriptomics analysis

Gene expression (GEX) analysis was conducted using Seurat (v5.0.1). Hashtag oligos were demultiplexed using the MULTIseqDemux function to assign sequences to the corresponding animals. Quality control measures excluded cells with fewer than 200 or more than 4,500 features, as well as those with mitochondrial RNA percentages exceeding 5%. Doublets and non-B cells were removed from the analysis. Datasets were then integrated using the anchor-based canonical correlation analysis (CCA) method. Further, Ig (LOC-) and MHC (MAMU-) genes were ignored for clustering.

### Statistical Analysis

Statistical analyses were conducted using Prism v10 (GraphPad Software). In the NHP study, the primary comparison was between group 1 and group 2 to evaluate differences between soluble versus membrane-bound MD39.3 mRNA immunogens. For pairwise comparisons between two groups at a single timepoint, the Mann-Whitney test was used. For pairwise comparisons between two groups across multiple timepoints, the Mann-Whitney test with Holm-Šídák correction for multiple comparisons was applied. For comparisons among multiple groups, the Kruskal-Wallis test with Dunn’s test for multiple comparisons was used.

## Acknowledgments

We thank D. Magnani via NHPRR for sharing purified anti-CD38 (OKT10) antibody. We thank R. Wilson for sharing 17b IgG. We thank A. Grifoni and P. Rubiro for lyophilizing the MD39 peptide pools. We are grateful to the Flow Cytometry and Next Generation Sequencing core facilities at La Jolla Institute for Immunology for their services. MD39.3 mRNA immunogens are currently being investigated in humans HVTN302 (Identifier: NCT05217641).

## Author contributions

P.R.R. designed, conducted, and analyzed B cell assays and sequencing. C.A.C. designed, conducted, and analyzed polyclonal and monoclonal antibody assays. E.M.Z. designed, conducted, and analyzed T cell assays, and assisted in GEX analysis. A.L. conducted ELISA assays. E.L., J.H.L., C.F., K.M., E.S., X.Z. carried out neutralization assays. J.L.T., A.M.J., W.H.L., and A.S.T. conducted EMPEM assays. A.M. assisted in carrying out the B cell sequencing. O.K. conducted SPR assays. E.G., N.P., D.L., S.E., M.K., and N.A. produced purified proteins. S.M. and B.G. supervised protein production, coordinated mRNA immunogens, and coordinated the rabbit study. S.B. and J.K.D. carried out, and J.R.Y. and J.C.P. supervised, mass spectrometry glycan profiling. T.S. and J.M.S. contributed to immunogen design. C.A.E. and V.L. provided assistance with the NHP study. A.P. and S.P.K. performed and analyzed bone marrow plasma cell responses. D.G.C. and G.S. supervised the NHP study. E.B.A., K.K.M. and D.J.I. provided SMNP. S.H. provided mRNA immunogens. D.S. supervised neutralization assays. G.O. and A.B.W. supervised EMPEM studies. S.C. and W.R.S. conceived and supervised the study. P.R.R., C.A.C., S.C. and W.R.S. wrote the paper. All authors reviewed and edited the paper.

## Data and materials availability

All data associated with this study are included in the paper or the Supplementary Materials. Requests for materials should be directed to the corresponding author, and a material transfer agreement (MTA) may apply. The scRNA-seq data have been deposited in the GEO and SRA databases under accession numbers GSE00000000 and SRX00000000, respectively.

---

**Fig. S1. Characterization of BG505 MD39 soluble immunogens.** (**A**) Biolayer interferometry (BLI) was used to assess antigenic profiles for the indicated trimers binding to IgGs for bnAbs (quaternary, PGT151 and PGT145; CD4bs, VRC01; and V3-glycan, PGT121 and PGT128) and non-nAbs (V3, 19b; CD4bs, B6 and F105). (**B**) Yield and thermostability of BG505 MD39 based immunogens. Yield was determined after 2G12 affinity chromatography and SEC purification. Thermostability measurements were made using nano differential scanning fluorimetry. (**C**) Negative stain electron microscopy analysis of BG505 MD39.3 gp140. (**D**) Glycan analysis of BG505 MD39 gp140 and BG505 MD39.3 gp140. Green indicates high mannose glycans, pink indicates complex type glycans and gray indicates unoccupied glycosylation sites. The N241 and N289 glycosylation sites are not present on BG505 MD39 gp140. Glycosylation sites are numbered using HxB2 numbering. *Adapted with permission from (*47*). (**E**) Negative stain electron microscopy analysis of BG505 SOSIP gp140 or BG505 MD39 gp140 with human sCD4. (**F**) Binding of sCD4 followed by 17b IgG for BG505 SOSIP, BG505 MD39, BG505 MD39.3, and BG505 MD39.3-CD4KO assessed by BLI.

**Fig. S2. Cell surface antigenicity.** Flow cytometry analysis of HEK293F cells transfected with DNA plasmids encoding membrane-bound HIV Env constructs. Cells were stained with bnAbs quaternary specific (PGT145 and PGT151), CD4bs (VRC01), V3-glycan (PGT121 and PGT128), MPER (10E8), or non-neutralizing antibodies (F105, B6, and 19b). (**A**) Raw MFI values show expression levels of each construct. Median values plotted with error bars showing the range (n=2). (**B**) PGT121 normalized MFI values showing antigenicity scaled to expression levels for each construct. (**C**) Comparison of effect of glycan hole for autologous neutralization. Week 26 serum neutralization against BG505 T332N pseudovirus of MD39.3 gp140 protein (G5) with glycan hole-containing MD39 gp140 protein NHP animals used in Silva et al (*20*). Bars indicate median for neutralization data. Statistical significance was assessed using the Mann-Whitney test. Significance levels were indicated as \**P* < 0.05, \*\**P* < 0.01, \*\*\**P* < 0.001, and \*\*\*\**P* < 0.0001.

**Fig. S3. Gating strategy for B_GC_ cells in LNs and B_Mem_ cells in PBMCs in NHPs.** (**A**) Flow cytometry gating strategy of Env-binding B_GC_ cells in LN FNAs. (**B**) Frequency of Env-binding B_GC_ cells in sampled LNs across groups and timepoints. (**C**) Frequency of total B_GC_ cells in sampled LNs across groups and timepoints. (**D**) Flow cytometry gating strategy of Env-binding B_Mem_ cells in PBMCs. Each point indicates a single animal (n=6/group) in (B) and the curves indicate the median. Each point indicates the mean with corresponding error in (C).

**Fig. S4. AIM/ICS T cell characterization and gating strategy.** (**A**) Flow cytometry gating strategy of AIM-ICS assay to detect antigen-specific CD4 T cells. (**B**) Quantification of Env-specific cytokine-producing (AIM^+^ICS^+^) CD4 T cells as a percentage of CD4 T cells for G2 and G5 at week 2 (post-prime) and week 26 (post-third). (**C**) Representative flow plots and quantification of Env-specific circulating T_FH_ cells as a percentage of CD4 T cells for G2 and G5 at week 2 (post-prime) and week 26 (post-third). Data are shown as background subtracted. Non-responder samples are set at baseline. The dotted black line indicates the limit of quantification (LOQ). Bars represent geometric mean and each point indicate a single animal (n=6/group). Animals with missing or poor-quality samples were excluded from the analysis. Statistical significance was assessed using the Kruskal-Wallis test, followed by Dunn’s multiple comparisons test. Significance levels were indicated as \**P* < 0.05, \*\**P* < 0.01, \*\*\**P* < 0.001, and \*\*\*\**P* < 0.0001.

**Fig. S5. mAb characterization of Env-binding B_Mem_ cells in NHPs.** (**A**) SPR data of mAb affinities and binding to different MD39.3 proteins with mAbs targeting the base epitope in green, the C3/V5 epitope in blue, and the V1/V3 epitope in orange. mAbs targeting unknown epitopes are shown in black. (**B**) Neutralization of mAbs tested for a panel of different viruses with the same coloring as (A). (**C**) Binding of mAbs to Link14 and V3 peptides assessed by BLI with the same coloring as (A).

**Fig. S6. V gene usage of Env-binding B_Mem_ cells in NHPs.** (**A**) V gene usage of heavy and light (kappa and lambda) chains of sequenced Env-binding B_Mem_ cells from G2 and G5. (**B**) CDRH3 length of sequenced Env-binding B_Mem_ cells from G2 and G5.

**Fig. S7. Transcriptomic analysis of Env-binding B_Mem_ cells in NHPs. (A**) Uniform manifold approximation and projection (UMAP) visualization of single-cell gene expression profiles identifying clusters among Env-binding B_Mem_ cells sorted from PBMCs. UMAP visualization split by week (week 8, 10, 24, 26 and 32) or by the number of nucleotide heavy-chain mutations (more than 30 mutations, between 30 to 15 mutations or less than 15 mutations). UMAP visualization depicting B cell isotypes in different colors. Feature plots showing the expression of *MS4A1* (CD20), *CD79B*, *JCHAIN*, and *MKI67*. (**B**) Heatmap displaying the 20 most upregulated genes for each cluster.

