## Supplemental Materials for "Vaccination with mRNA-encoded membrane-bound HIV Envelope trimer induces neutralizing antibodies in animal models"

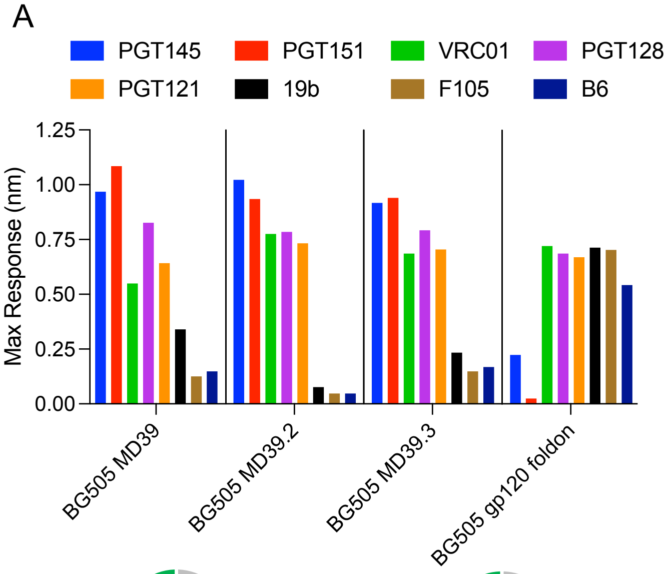


B

| Name | Post SEC Yield (mg/L) | Tm (°C) |
|---|---|---|
| BG505 MD39 gp140 | 3.6 | 78.7 |
| BG505 MD39.2 gp140 | 3.3 | 73.6 |
| BG505 MD39.3 gp140 | 3.1 | 75.6 |

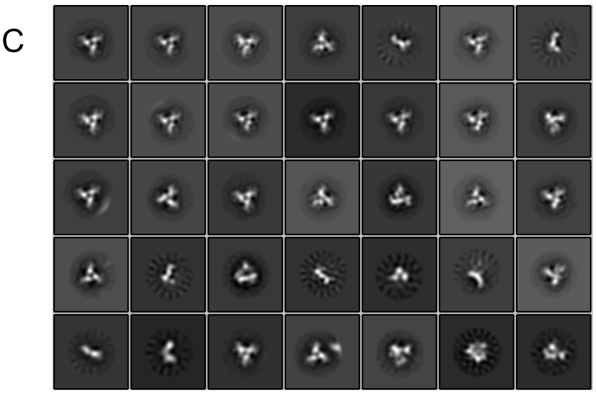


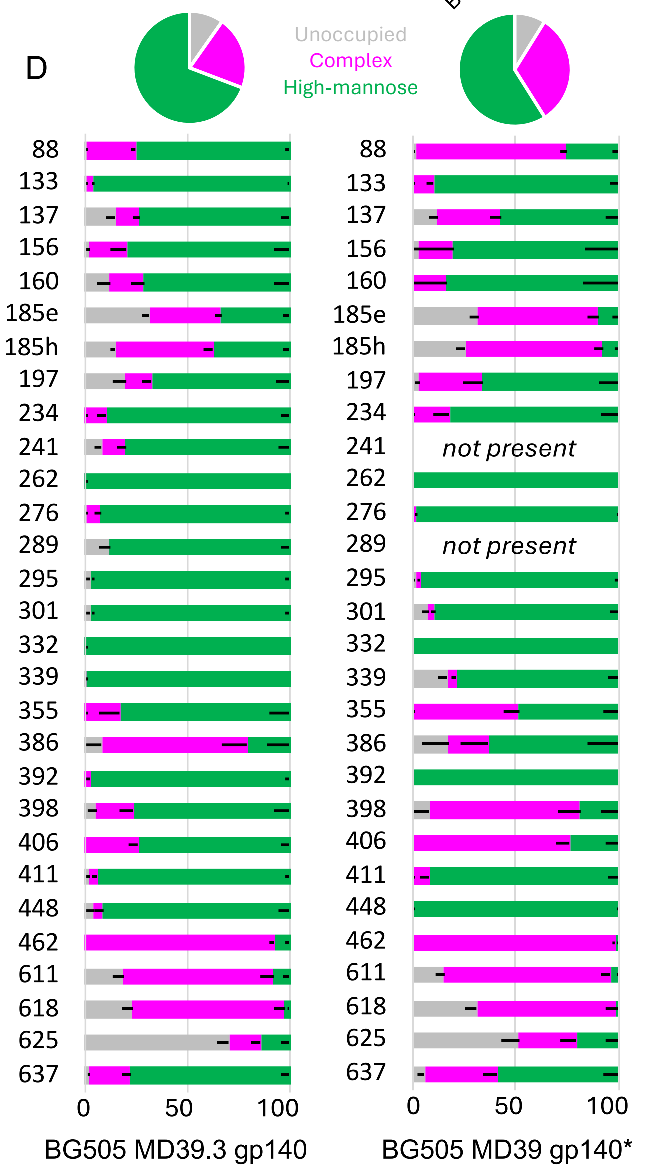


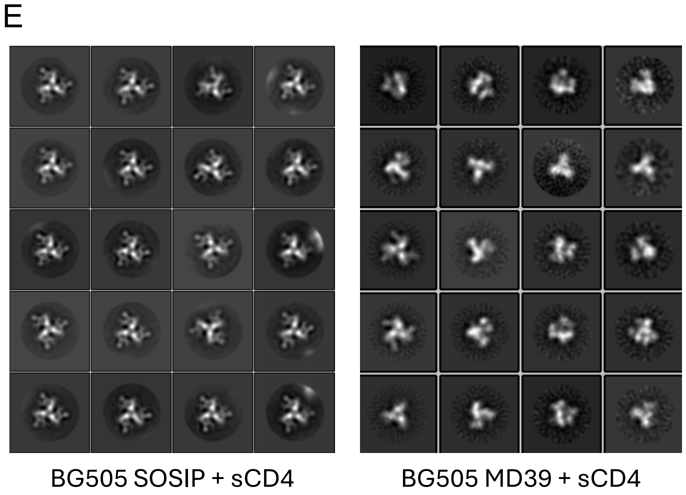


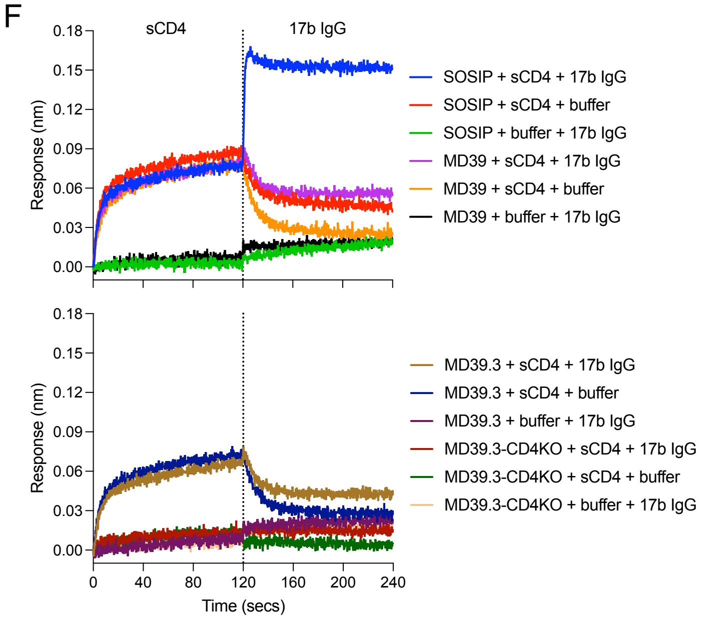


Figure S1

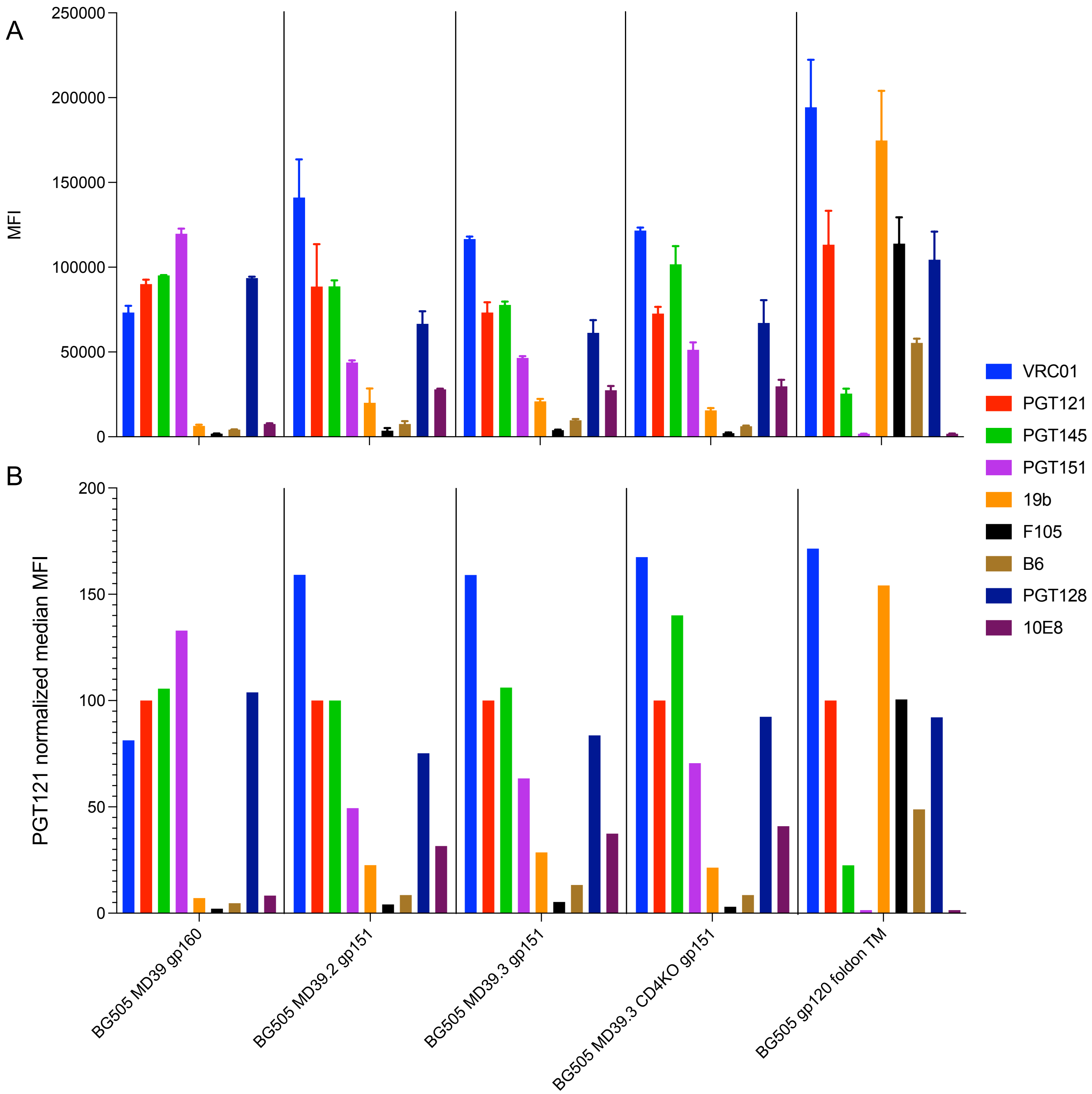


C

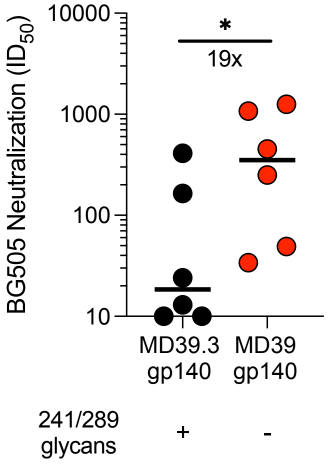


| group | animal | ID50 | geo mean | median |
|---|---|---|---|---|
| MD39.3 gp140 | RBa19 | 13 | 36 | 19 |
| | REy18 | 165 | | |
| | RWh18 | 10 | | |
| | RWm18 | 410 | | |
| | RWn18 | 24 | | |
| | RYt17 | 10 | | |
| MD39 gp140 | RJc18 | 251 | 252 | 353 |
| | RMs17 | 34 | | |
| | RGu17 | 454 | | |
| | 34880 | 49 | | |
| | RNv17 | 1252 | | |
| | RUs17 | 1076 | | |

Figure S2

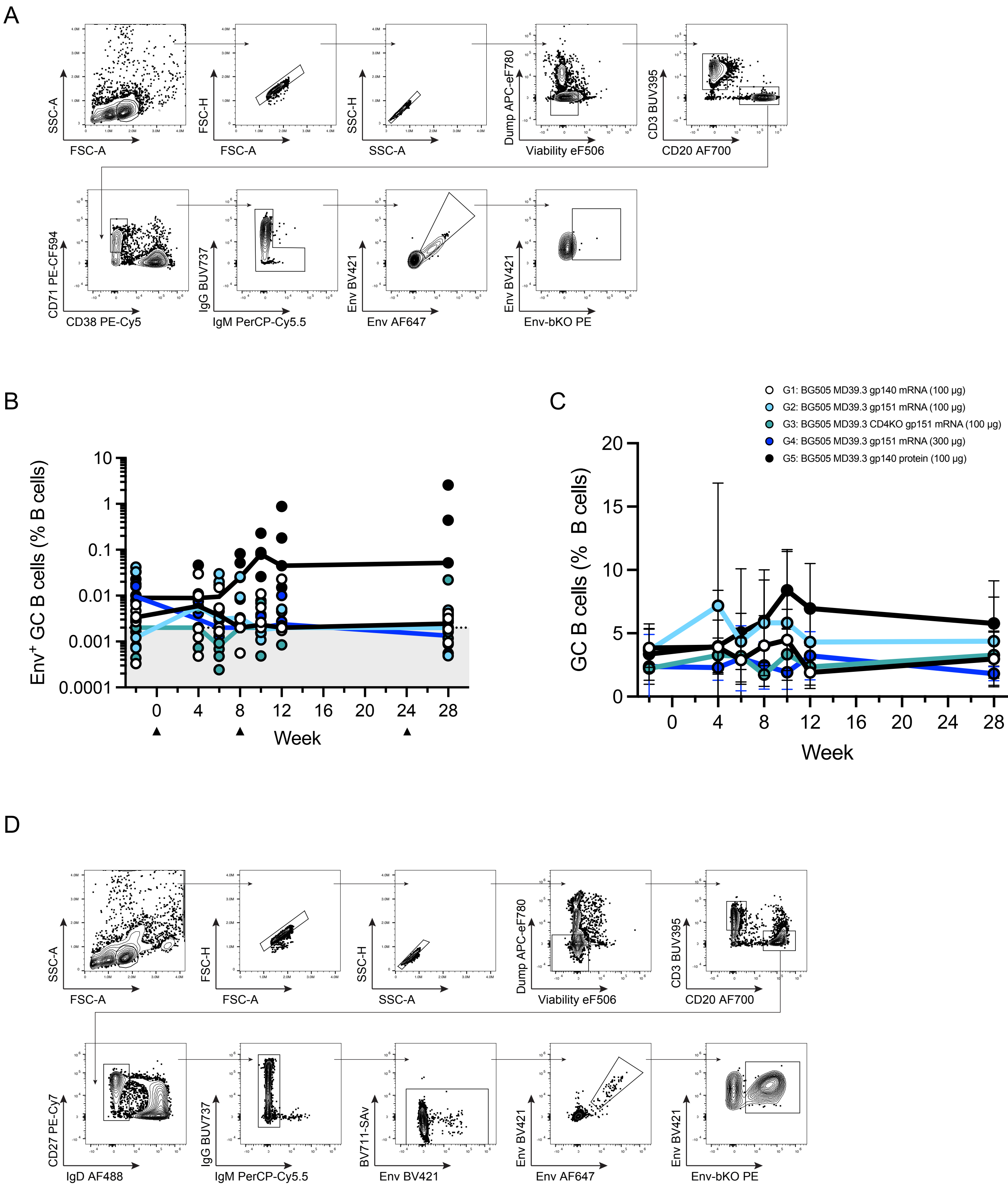
A
SSC-A
FSC-A
FSC-H
FSC-A
SSC-H
SSC-A
Dump APC-eF780
Viability eF506
CD3 BUV395
CD20 AF700
CD71 PE-CF594
CD38 PE-Cy5
IgG BUV737
IgM PerCP-Cy5.5
Env BV421
Env AF647
Env BV421
Env-bKO PE
B
Env+ GC B cells (% B cells)
10
1
0.1
0.01
0.001
0.0001
0 4 8 12 16 20 24 28
Week
C
G1: BG505 MD39.3 gp140 mRNA (100 µg)
G2: BG505 MD39.3 gp151 mRNA (100 µg)
G3: BG505 MD39.3 CD4KO gp151 mRNA (100 µg)
G4: BG505 MD39.3 gp151 mRNA (300 µg)
G5: BG505 MD39.3 gp140 protein (100 µg)
GC B cells (% B cells)
20
15
10
5
0
0 4 8 12 16 20 24 28
Week
D
SSC-A
FSC-A
FSC-H
FSC-A
SSC-H
SSC-A
Dump APC-eF780
Viability eF506
CD3 BUV395
CD20 AF700
CD27 PE-Cy7
IgD AF488
IgG BUV737
IgM PerCP-Cy5.5
BV711-SAv
Env BV421
Env BV421
Env AF647
Env BV421
Env-bKO PE

Figure S3

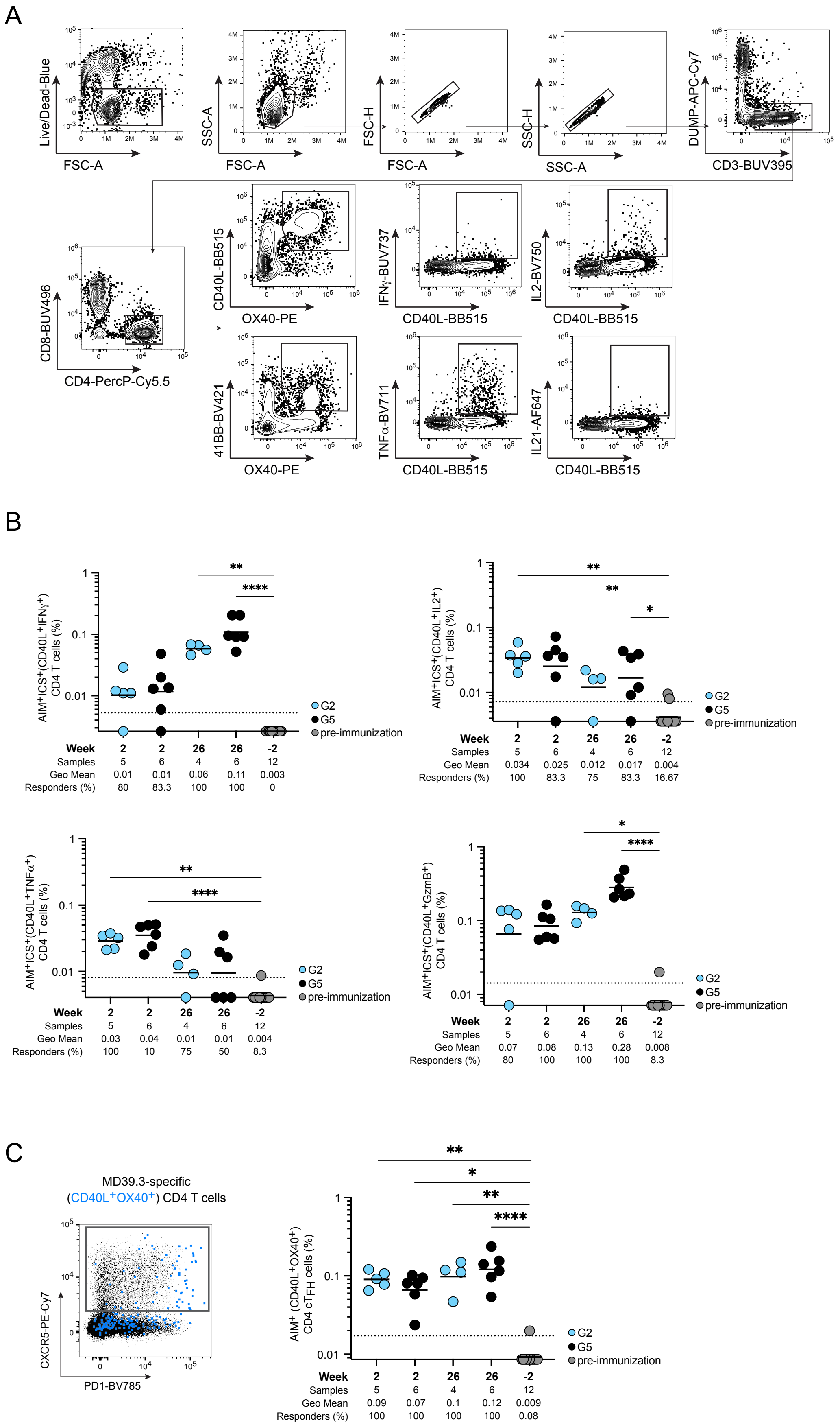

A
Live/Dead-Blue
FSC-A
SSC-A
FSC-A
FSC-H
FSC-A
SSC-H
SSC-A
DUMP-APC-Cy7
CD3-BUV395
CD8-BUV496
CD4-PercP-Cy5.5
CD40L-BB515
OX40-PE
IFNγ-BUV737
CD40L-BB515
IL2-BV750
CD40L-BB515
41BB-BV421
OX40-PE
TNFα-BV711
CD40L-BB515
IL21-AF647
CD40L-BB515
B
AIM+ICS+(CD40L+IFNγ+) CD4 T cells (%)
G2
G5
pre-immunization
Week 2 2 26 26 -2
Samples 5 6 4 6 12
Geo Mean 0.01 0.01 0.06 0.11 0.003
Responders (%) 80 83.3 100 100 0
AIM+ICS+(CD40L+IL2+) CD4 T cells (%)
Week 2 2 26 26 -2
Samples 5 6 4 6 12
Geo Mean 0.034 0.025 0.012 0.017 0.004
Responders (%) 100 83.3 75 83.3 16.67
AIM+ICS+(CD40L+TNFα+) CD4 T cells (%)
Week 2 2 26 26 -2
Samples 5 6 4 6 12
Geo Mean 0.03 0.04 0.01 0.01 0.004
Responders (%) 100 10 75 50 8.3
AIM+ICS+(CD40L+GzmB+) CD4 T cells (%)
Week 2 2 26 26 -2
Samples 5 6 4 6 12
Geo Mean 0.07 0.08 0.13 0.28 0.008
Responders (%) 80 100 100 100 8.3
C
MD39.3-specific
(CD40L+OX40+) CD4 T cells
CXCR5-PE-Cy7
PD1-BV785
AIM+ (CD40L+OX40+) CD4 cTFH cells (%)
G2
G5
pre-immunization
Week 2 2 26 26 -2
Samples 5 6 4 6 12
Geo Mean 0.09 0.07 0.1 0.12 0.009
Responders (%) 100 100 100 100 0.08


Figure S4

A

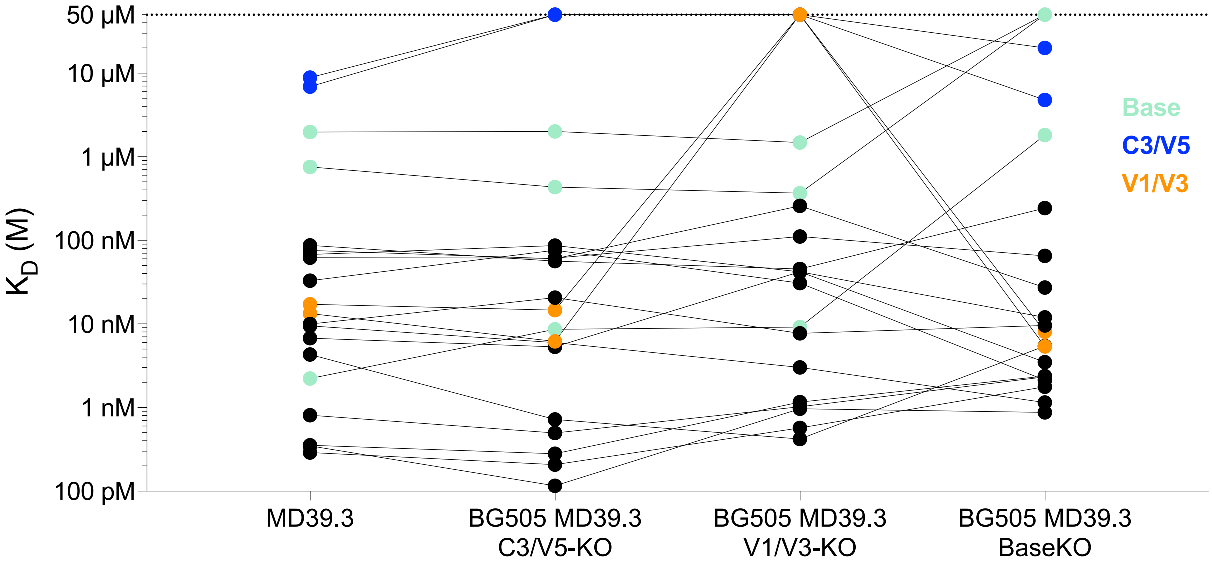


B

| | BG505 WT | BG505 T332N | BG505 N611A | BG505 N241+N289 | BG505 133aN | BG505 133aN+136aA | BG505 T465N | MLV |
|---|---|---|---|---|---|---|---|---|
| VRC01 | 0.084 | 0.047 | 0.089 | 0.050 | 0.120 | 0.091 | 0.084 | NN |
| RFu18_1_IgG | NN | NN | NN | NN | NN | NN | NN | NN |
| RFu18_10_IgG | NN | NN | NN | NN | NN | NN | NN | NN |
| RFu18_12_IgG | NN | NN | NN | NN | NN | NN | NN | NN |
| RFu18_2_IgG | NN | NN | NN | NN | NN | NN | NN | NN |
| RFu18_4_IgG | NN | NN | NN | NN | NN | NN | NN | NN |
| RFu18_6_IgG | NN | NN | NN | NN | NN | NN | NN | NN |
| RFu18_37_IgG | NN | NN | NN | NN | NN | NN | NN | NN |
| RFu18_38_IgG | NN | NN | NN | NN | NN | NN | NN | NN |
| RUv18_29_IgG | NN | NN | NN | NN | NN | NN | NN | NN |
| RUv18_30_IgG | NN | NN | NN | NN | NN | NN | NN | NN |
| RUv18_31_IgG | NN | NN | NN | NN | NN | NN | NN | NN |
| RUv18_40_IgG | NN | NN | NN | NN | NN | NN | NN | NN |
| RUv18_42_IgG | NN | NN | NN | NN | NN | NN | NN | NN |
| RUv18_43_IgG | 1.477 | 4.154 | 1.072 | 12.911 | NN | NN | 13.557 | NN |
| RFu18_11_IgG | NN | 21.955 | NN | NN | NN | NN | NN | NN |
| RUv18_44_IgG | 0.094 | 0.549 | 0.360 | 0.730 | NN | NN | 0.960 | NN |
| RUv18_46_IgG | NN | NN | NN | NN | NN | NN | NN | NN |
| RUv18_47_IgG | NN | NN | NN | NN | NN | NN | NN | NN |
| RWo17_32_IgG | NN | NN | NN | NN | NN | NN | NN | NN |
| DEN | NN | NN | NN | NN | NN | NN | NN | NN |

| $IC_{50}$ (μg/mL) |
|---|
| 0.001-0.010 |
| 0.010-0.100 |
| 0.100-1.000 |
| 1.000-10.00 |
| 10.00-50.00 |

C

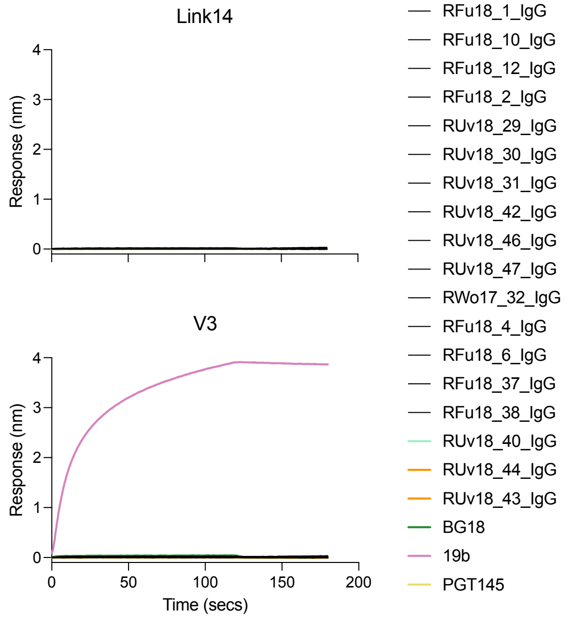


Figure S5

A

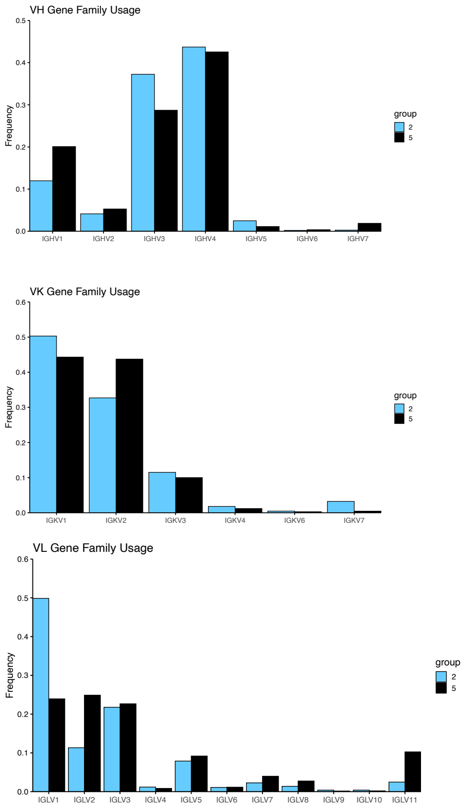

VH Gene Family Usage
Frequency
0.5
0.4
0.3
0.2
0.1
0.0
IGHV1
IGHV2
IGHV3
IGHV4
IGHV5
IGHV6
IGHV7
group
2
5
VK Gene Family Usage
Frequency
0.6
0.5
0.4
0.3
0.2
0.1
0.0
IGKV1
IGKV2
IGKV3
IGKV4
IGKV6
IGKV7
group
2
5
VL Gene Family Usage
Frequency
0.6
0.5
0.4
0.3
0.2
0.1
0.0
IGLV1
IGLV2
IGLV3
IGLV4
IGLV5
IGLV6
IGLV7
IGLV8
IGLV9
IGLV10
IGLV11
group
2
5


B

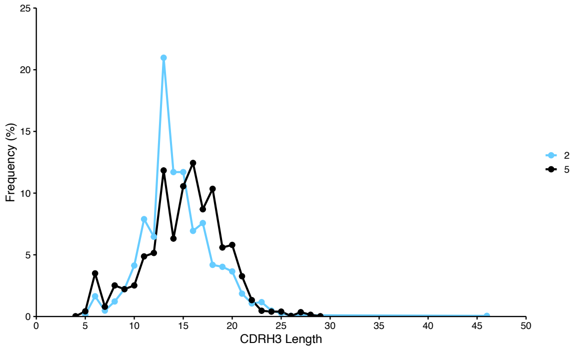

Frequency (%)
25
20
15
10
5
0
0
5
10
15
20
25
30
35
40
45
50
CDRH3 Length
2
5


Figure S6

A

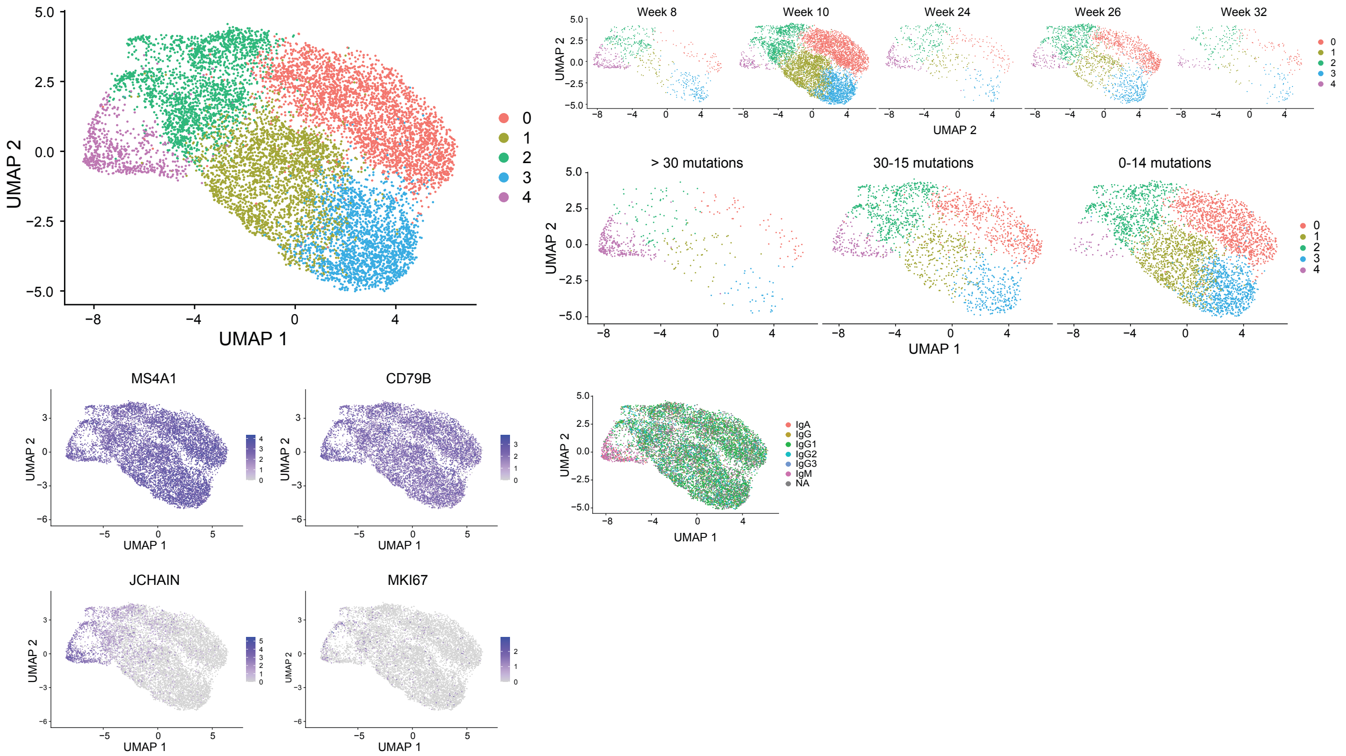

UMAP 1
UMAP 2
0
1
2
3
4
Week 8
Week 10
Week 24
Week 26
Week 32
> 30 mutations
30-15 mutations
0-14 mutations
MS4A1
CD79B
JCHAIN
MKI67
IgA
IgG
IgG1
IgG2
IgG3
IgM
NA


B

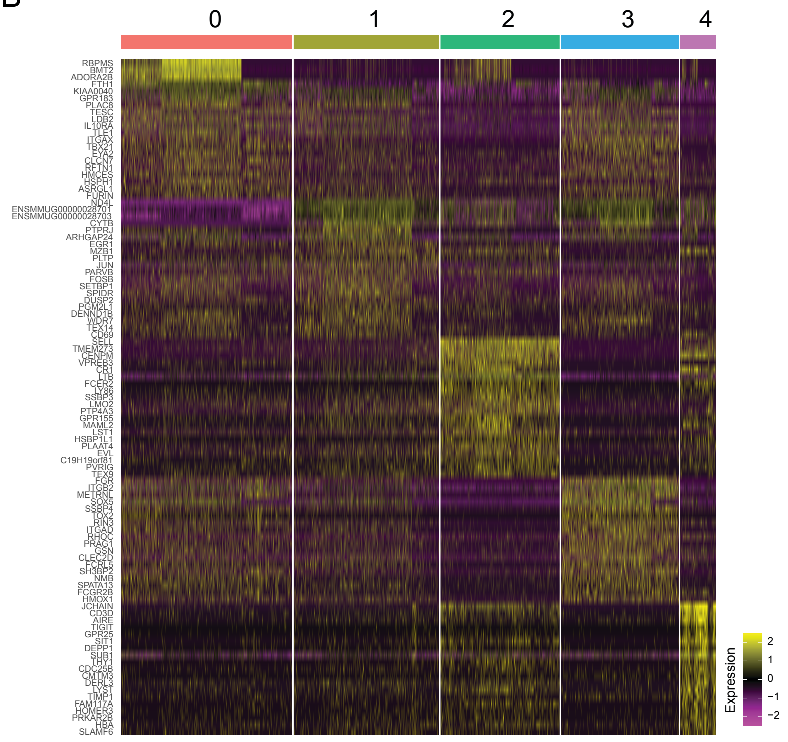

0
1
2
3
4
RBPMS
BMT2
ADORA2B
FTH1
KIAA0040
GPR183
PLAC8
TESC
LDB2
IL10RA
TLE1
ITGAX
TBX21
EYA2
CLCN7
RFTN1
HMCES
HSPH1
ASRGL1
FURIN
ND4L
ENSMMUG00000028701
ENSMMUG00000028703
CYTB
PTPRJ
ARHGAP24
EGR1
MZB1
PLTP
JUN
PARVB
FOSB
SETBP1
SPIDR
DUSP2
PGM2L1
DENND1B
WDR7
TEX14
CD69
SELL
TMEM273
CENPM
VPREB3
CR1
LTB
FCER2
LY86
SSBP3
LMO2
PTP4A3
GPR155
MAML2
LST1
HSBP1L1
PLAAT4
EVL
C19H19orf81
PVRIG
TEX9
FGR
ITGB2
METRNL
SOX5
SSBP4
TOX2
RIN3
ITGAD
RHOC
PRAG1
GSN
CLEC2D
FCRL5
SH3BP2
NMB
SPATA13
FCGR2B
HMOX1
JCHAIN
CD3D
AIRE
TIGIT
GPR25
SIT1
DEPP1
SUB1
THY1
CDC25B
CMTM3
DERL3
LYST
TIMP1
FAM117A
HOMER3
PRKAR2B
HBA
SLAMF6
Expression


Figure S7

**Table S1. Rabbit study schema.** All animals were immunized at weeks 0, 8, and 24.

| Group | N | Immunogen | mRNA/Protein | Dose ($\mu$g) | Route | Adjuvant |
|---|---|---|---|---|---|---|
| 1 | 6 | BG505 MD39.2 gp140 | mRNA | 100 | IM | none |
| 2 | 6 | BG505 MD39.3 gp140 | mRNA | 100 | IM | none |
| 3 | 6 | BG505 MD39.2 gp151 | mRNA | 100 | IM | none |
| 4 | 6 | BG505 MD39.3 gp151 | mRNA | 100 | IM | none |
| 5 | 6 | BG505 MD39.3 CD4KO gp151 | mRNA | 100 | IM | none |
| 6 | 6 | BG505 gp120 foldon | Protein | 30 | IM | SMNP (375 $\mu$g) |
| 7 | 6 | BG505 MD39.3 soluble trimer | Protein | 30 | IM | SMNP (375 $\mu$g) |
| 8 | 6 | BG505 MD39 soluble trimer | Protein | 30 | IM | SMNP (375 $\mu$g) |

**Table S2. Flow Panel for $B_{Mem}$ analysis**

| Antibodies | Clone | Source | Cat # |
|---|---|---|---|
| Streptavidin BV711 (empty) | - | BioLegend | 405241 |
| Streptavidin AF647 (MD39.3) | - | BioLegend | 405237 |
| Streptavidin BV421 (MD39.3) | - | BioLegend | 405225 |
| Streptavidin PE (MD39.3 bKO) | - | BioLegend | 405245 |
| Fixable Viability Dye eFluor506 | - | Thermo Fisher Scientific | 65-0866-18 |
| Mouse anti-human CD4 BV650 | OKT4 | BioLegend | 317436 |
| Mouse anti-human CD8a APC-eFluor780 | RPA-T8 | Thermo Fisher Scientific | 47-0088-42 |
| Mouse anti-human CD16 APC-eFluor780 | CB16 | Thermo Fisher Scientific | 47-0168-42 |
| Mouse anti-human CD20 AF700 | 2H7 | BioLegend | 302322 |
| Mouse anti-human IgG BUV737 | G18-145 | BD Biosciences | 612819 |
| Mouse anti-human CD27 PE-Cy7 | O323 | Thermo Fisher Scientific | 25-0279-42 |
| Mouse anti-human CD3 BUV395 | SP34-2 | BD Biosciences | 564117 |
| Goat anti-human IgD AF488 | polyclonal | SouthernBiotech | 2030-30 |
| Mouse anti-human IgM PerCP-Cy5.5 | G20-127 | BD Biosciences | 561285 |

**Table S3. Flow Panel for $B_{GC}$ analysis**

| Antibodies | Clone | Source | Cat # |
|---|---|---|---|
| Streptavidin AF647 (MD39.3) | - | BioLegend | 405237 |
| Streptavidin BV421 (MD39.3) | - | BioLegend | 405225 |
| Streptavidin PE (MD39.3 bKO) | - | BioLegend | 405245 |
| Fixable Viability Dye eFluor506 | - | Thermo Fisher Scientific | 65-0866-18 |
| Mouse anti-human CD4 BV711 | OKT4 | BioLegend | 317440 |
| Mouse anti-human CD8a APC-eFluor780 | RPA-T8 | Thermo Fisher Scientific | 47-0088-42 |
| Mouse anti-human CD16 APC-eFluor780 | CB16 | Thermo Fisher Scientific | 47-0168-42 |
| Mouse anti-human CD20 AF488 | 2H7 | BioLegend | 302316 |
| Mouse anti-human IgG BUV737 | G18-145 | BD Biosciences | 612819 |
| Mouse anti-human CXCR5 PE-Cy7 | Mu5UBEE | Thermo Fisher Scientific | 25-9185-42 |
| Mouse anti-human CD3 BUV395 | SP34-2 | BD Biosciences | 564117 |
| Mouse anti-rhesus CD38 PE-Cy5 | OKT10 | In house | - |
| Mouse anti-human IgM PerCP-Cy5.5 | G20-127 | BD Biosciences | 561285 |
| Mouse anti-human PD1 BV605 | EH12.2H7 | BioLegend | 329924 |
| Mouse anti-human CD71 PE-CF594 | L01.1 | BD Biosciences | custom |

**Table S4. Flow Panel for sorting of MD39.3-specific $B_{Mem}$ cells**

| Antibodies | Clone | Source | Cat # |
|---|---|---|---|
| Streptavidin AF647 (MD39.3) | - | BioLegend | 405237 |
| TotalSeq-C Streptavidin BV421 (MD39.3) | - | BioLegend | custom |
| TotalSeq-C Streptavidin PE (MD39.3 bKO) | - | BioLegend | 405155 |
| Fixable Viability Dye eFluor506 | - | Thermo Fisher Scientific | 65-0866-18 |
| Mouse anti-human CD8a APC-eFluor780 | RPA-T8 | Thermo Fisher Scientific | 47-0088-42 |
| Mouse anti-human CD16 APC-eFluor780 | CB16 | Thermo Fisher Scientific | 47-0168-42 |
| Mouse anti-human CD14 APC-Cy7 | M5E2 | BioLegend | 301820 |
| Mouse anti-human CD3 APC-Cy7 | SP34-2 | BD Biosciences | 557757 |
| Mouse anti-human CD20 BUV395 | 2H7 | BD Biosciences | 563781 |
| Mouse anti-human IgG BV605 | G18-145 | BD Biosciences | 563246 |
| Mouse anti-human CD27 PE-Cy7 | O323 | Thermo Fisher Scientific | 25-0279-42 |
| Goat anti-human IgD AF488 | polyclonal | SouthernBiotech | 2030-30 |
| Mouse anti-human IgM PerCP-Cy5.5 | G20-127 | BD Biosciences | 561285 |

**Table S5. Flow cytometry AIM and ICS panel staining for MD39.3-specific T cells**

| Antibodies | Clone | Source | Cat # |
|---|---|---|---|
| LIVE/DEAD Fixable Blue | - | Invitrogen | L23105 |
| GolgiPlug | - | BD Biosciences | 555029 |
| GolgiStop | - | BD Biosciences | 554724 |
| Mouse anti-human CD40 | HB14 | Miltenyi | 130-094-133 |
| Mouse anti-human CXCR5 PE-Cy7 | MU5UBEE | Thermo Fisher Scientific | 25-9185-42 |
| Mouse anti-human CCR7 BV650 | G043H7 | BioLegend | 353233 |
| Mouse anti-human CD69 PE-Cy5 | FN50 | BioLegend | 310908 |
| Mouse anti-human CD137 (4-1BB) BV421 | 4B4-1 | BioLegend | 309819 |
| Mouse anti-human CD25 BV605 | BC96 | BioLegend | 302631 |
| Mouse anti-human CD40L BB515 | 24-31 | BD Biosciences | 568170 |
| Mouse anti-human CD134 (OX40) PE | L106 | BD Biosciences | 340420 |
| Mouse anti-human CD8 BUV496 | RPA-T8 | BD Biosciences | 612943 |
| Mouse anti-human CD14 APC-Cy7 | M5E2 | BioLegend | 301820 |
| Mouse anti-human CD16 APC-eFluor780 | eBioCB16 | Thermo Fisher Scientific | 47-0168-42 |
| Mouse anti-human CD20 APC-Cy7 | 2H7 | BioLegend | 302314 |
| Mouse anti-human CD3 BUV395 | SP34-2 | BD Biosciences | 564117 |
| Mouse anti-human CD4 PerCP-Cy5.5 | OKT4 | BioLegend | 317428 |
| Mouse anti-human PD-1 BV785 | EH12.2H7 | BioLegend | 329929 |
| Mouse anti-human CD45RA PE-CF594 | 5H9 | BD Biosciences | 565419 |
| Armenian Hamster anti-ICOS BV480 | C398.4A | BD Biosciences | 566087 |
| Mouse anti-human IFN-γ BUV737 | 4S.B3 | BD Biosciences | 612845 |
| Rat anti-human IL-2 BV750 | MQ1-17H12 | BD Biosciences | 566361 |
